# Multivalent designed proteins protect against SARS-CoV-2 variants of concern

**DOI:** 10.1101/2021.07.07.451375

**Authors:** Andrew C. Hunt, James Brett Case, Young-Jun Park, Longxing Cao, Kejia Wu, Alexandra C. Walls, Zhuoming Liu, John E. Bowen, Hsien-Wei Yeh, Shally Saini, Louisa Helms, Yan Ting Zhao, Tien-Ying Hsiang, Tyler N. Starr, Inna Goreshnik, Lisa Kozodoy, Lauren Carter, Rashmi Ravichandran, Lydia B. Green, Wadim L. Matochko, Christy A. Thomson, Bastain Vögeli, Antje Krüger-Gericke, Laura A. VanBlargan, Rita E. Chen, Baoling Ying, Adam L. Bailey, Natasha M. Kafai, Scott Boyken, Ajasja Ljubetič, Natasha Edman, George Ueda, Cameron Chow, Amin Addetia, Nuttada Panpradist, Michael Gale, Benjamin S. Freedman, Barry R. Lutz, Jesse D. Bloom, Hannele Ruohola-Baker, Sean P. J. Whelan, Lance Stewart, Michael S. Diamond, David Veesler, Michael C. Jewett, David Baker

## Abstract

Escape variants of SARS-CoV-2 are threatening to prolong the COVID-19 pandemic. To address this challenge, we developed multivalent protein-based minibinders as potential prophylactic and therapeutic agents. Homotrimers of single minibinders and fusions of three distinct minibinders were designed to geometrically match the SARS-CoV-2 spike (S) trimer architecture and were optimized by cell-free expression and found to exhibit virtually no measurable dissociation upon binding. Cryo-electron microscopy (cryoEM) showed that these trivalent minibinders engage all three receptor binding domains on a single S trimer. The top candidates neutralize SARS-CoV-2 variants of concern with IC_50_ values in the low pM range, resist viral escape, and provide protection in highly vulnerable human ACE2-expressing transgenic mice, both prophylactically and therapeutically. Our integrated workflow promises to accelerate the design of mutationally resilient therapeutics for pandemic preparedness.

**One-Sentence Summary:** We designed, developed, and characterized potent, trivalent miniprotein binders that provide prophylactic and therapeutic protection against emerging SARS-CoV-2 variants of concern.

## Main Text

Monoclonal antibodies (mAbs) targeting the SARS-CoV-2 spike (S) glycoprotein can improve disease outcomes for patients with COVID-19. However, producing mAbs in sufficient quantities for population scale use during a global pandemic is technically and financially challenging (*1*), and many mAbs are sensitive to viral escape via point mutations in their recognition epitope on the S trimer (*2, 3*). To overcome this limitation, it is common practice to prepare a cocktail of different mAbs targeting different epitopes. However, two circulating SARS-CoV-2 variants, B.1.351 (Beta) and P.1 (Gamma), disrupt binding of both mAbs in the authorized bamlanivimab and etesevimab cocktail as well as casirivimab in the authorized REGN-COV cocktail (*3–6*). Furthermore, in polyclonal sera elicited by the authorized COVID-19 mRNA vaccines, a small number of point mutations cause significant reductions in neutralization capacity (*2, 7–10*). As a result, the rapidly spreading variants, B.1.1.7 (Alpha), B.1.351 (Beta), P.1 (Gamma), and B.1.617.2 (Delta), have raised significant concern about the possibility for escape from currently authorized vaccines and therapeutics. Together with the slow rollout of vaccines globally, this highlights the urgent need for prophylactic and therapeutic interventions whose efficacy is not disrupted by the ongoing antigenic drift, as is the case for a few mAbs (*11–18*).

As an alternative to mAbs, we previously computationally designed miniproteins that block the SARS-CoV-2 receptor binding domain (RBD) interaction of the S trimer with its host receptor ACE2 (*19*). An ACE2-mimic, AHB2, which incorporates the primary ACE2-RBD-interacting helix in a custom designed small 3-helix bundle, and two *de novo* designs, LCB1 and LCB3, with new RBD binding interfaces, neutralize the Wuhan-1 SARS-CoV-2 virus with IC_50_ values in the pM to nM range. LCB1 has protective activity as both a pre-exposure prophylactic and post-exposure therapeutic in human ACE2 (hACE2) transgenic mice (*20*). The designs are expressed at high levels in *Escherichia coli* and are highly thermostable, requiring only heat treatment followed by ion-exchange chromatography to achieve high purity (Fig. S1), which could considerably streamline manufacturing and decrease the cost of goods. To determine the potential for mutations to arise that disrupt LCB1 and AHB2 binding to the RBD, we performed deep mutational scans using site saturation mutagenesis of the RBD. We found that for LCB1, the widely observed K417N mutation results in a likely greater than 10-fold reduction in affinity and the E406W and Y453K/R mutations result in a likely greater than 100-fold reduction in affinity, each without strongly reducing RBD-ACE2 affinity (Fig. S2). For AHB2, we similarly observed several mutations, including K417N, E406W, and Y453K/R that reduce the affinity of the minibinder for the RBD.

### Multivalent minibinder design and experimental optimization

To improve the ability of the minibinders to neutralize currently circulating SARS-CoV-2 variants, we developed multivalent versions of the minibinders with geometries enabling simultaneous engagement of all 3 RBDs in a single S trimer. We hypothesized that such constructs would substantially increase binding affinity through avidity by occupying several RBDs. Further, we reasoned this could enable the multivalent minibinders to be largely insensitive to mutations that would escape binding of the monovalent minibinders (a 100x reduction in binding affinity of a sub-picomolar binder would still result in an affinity in a therapeutic range in a multivalent construct). Additionally, we reasoned that constructs with binding domains engaging distinct epitopes or containing different sets of contacts with the target epitope could prevent escape. To design multivalent constructs, we started from optimized versions of the previously described LCB1, AHB2, and LCB3 minibinders (hereafter referred to as monomers MON1, MON2, and MON3, respectively; Table S1). To assess whether multivalency would improve the breadth of minibinders as a therapeutic for emerging variants of concern, we developed a cell-free protein synthesis (CFPS) workflow which combines an *in vitro* DNA assembly step followed by PCR to generate linear expression templates that are used to drive CFPS and enable rapid prototyping of new minibinder designs (Fig. S3). The workflow enables assembly and translation of synthetic genes and generation of purified protein in as little as 6 hours, is compatible with high-throughput, automated experimentation, and is easily scaled for the production of mg quantities of protein (*21, 22*). For evaluation, we coupled the workflow to an AlphaLISA protein-protein interaction (PPI) competition assay to enable comparison of dissociation rates of the designed proteins against either the monomeric RBD or the trimeric HexaPro SARS-CoV-2-S-glycoprotein (S6P) (*23*). Because multivalency largely impacts the dissociation rate constant of the protein-protein interaction, we reasoned that an in-solution off-rate screen could distinguish differences between mono and multivalent binding (*24*). Multivalent minibinders were allowed to fully associate with the target protein, then reactions were split in two and either 100-fold molar excess of untagged competitor (to prevent reassociation) or buffer was added. The ratio of the competitor to no-competitor condition measurements were calculated to determine the fraction of the complex dissociated (*25*).

Paralleling previous work where trimeric binders were targeted to the sialic acid binding site on influenza hemagglutinin (*26*), we first designed homotrimeric versions of the MON1, MON2, and MON3 miniproteins geometrically matched to the three RBDs in the S trimer (hereafter referred to as TRI; for example, TRI1-1 represents a homotrimer of MON1 with homotrimerization domain 1, Table S1). We designed and screened more than 100 different homotrimeric minibinders, with varied linker lengths and homotrimzeriation domains, using the CFPS workflow. We observed that many of the homotrimeric constructs exhibited slower dissociation rates than the corresponding monomers; much larger effects were observed with dissociation from the S trimer than monomeric RBD consistent with multivalent binding (Fig. 1 and Fig. S4C-E). The top binders exhibited little to no dissociation from S trimer after 7 days of incubation with competitor, indicating a likely apparent dissociation rate constant of 1×10^−7^ s^−1^ or slower. This is a marked increase, more than four orders of magnitude for the TRI2 proteins, over the dissociation rate constants of the monomeric minibinders (Fig. S5). We selected two trimeric scaffolds, the designed two ring helical bundle SB175 and the T4 foldon (*27*) (Table S2), to proceed with based on the screening results and previous experience with these scaffolds.

**Fig. 1.**
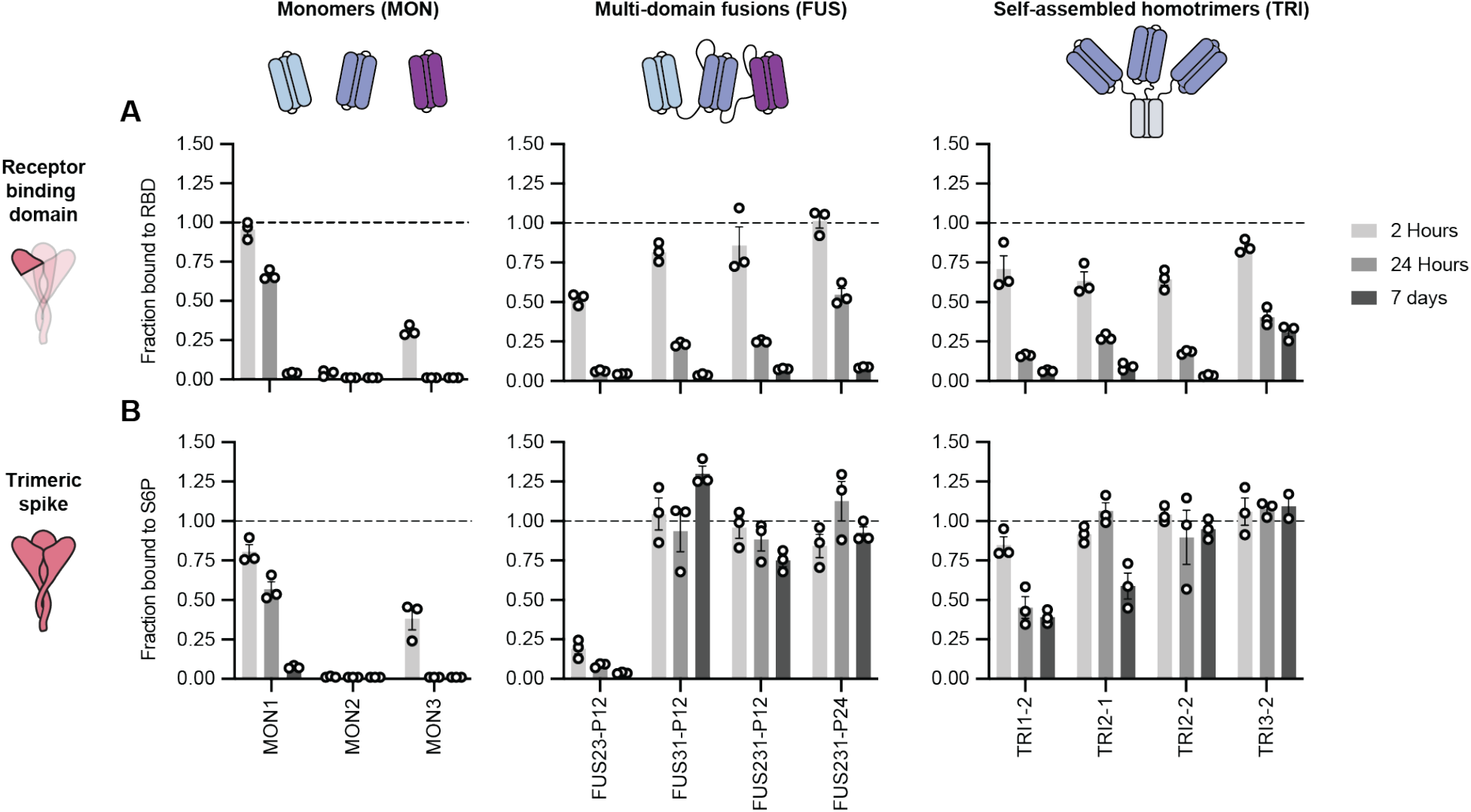
Multivalent minibinders exhibit unusually slow dissociation rates upon binding to the prefusion SARS-CoV-2-S glycoprotein. (**A, B**) Dissociation of the minibinder construct complexed with either the receptor binding domain (RBD) (**A**) or S trimer (S6P) (**B**) was monitored via competition with 100-fold molar excess of untagged MON1 using AlphaLISA (Mean ± SEM, n = 3 replicates from a single experiment).

Next, we generated two- and three-domain fusions of the MON1, MON2, and MON3 minibinders separated by flexible linkers (hereafter referred to as FUS; for example, FUS31-P12 represents a fusion of MON3 to MON1 separated by a 12 amino acid proline-alanine-serine (PAS12) linker, Table S1). We screened more than 100 different fusions using the CFPS workflow, evaluating different minibinder orderings and a range of linker compositions and lengths that span the distances between the termini of the domains when bound to the “open” and “closed” states of the RBD (Fig. 1 and Fig. S4A-B, and F) (*28*). FUS31 and FUS231 constructs showed slower dissociation against S6P than RBD, and exhibited slower dissociation than all monomeric minibinders tested, consistent with multivalent S6P engagement. The top binders exhibited little dissociation from S6P after 7 days, indicating a likely apparent dissociation rate constant of 1×10^−7^ s^−1^ or slower, representing, at minimum, more than one order of magnitude improvement over the strongest monomeric dissociation rate constant (Fig. S5).

### Structural studies of minibinders in complex with SARS-CoV-2 S

We next sought to determine the extent to which the designed multivalent constructs engage multiple RBDs on a single S trimer (on a virion, the S trimers are too far apart for single compounds to bind more than one, hence avidity requires multivalent engagement within a single trimer (*29–31*)). Initial screening revealed considerable cross-linking and aggregation of S proteins upon addition of constructs FUS31-G8 or TRI1-5 (Fig. S6), consistent with binding to RBDs on different S proteins. In contrast, for constructs TRI2-2, FUS231-G10, FUS231-P24 and FUS31-G10, we observed little cross-linking, consistent with multivalent engagement of a single S trimer for each minibinder. To determine the binding modes of these compounds to the S trimer and characterize the structure of the MON2 and RBD interactions at high resolution, we carried out cryoEM characterization of these complexes (Fig. 2).

**Fig. 2.**
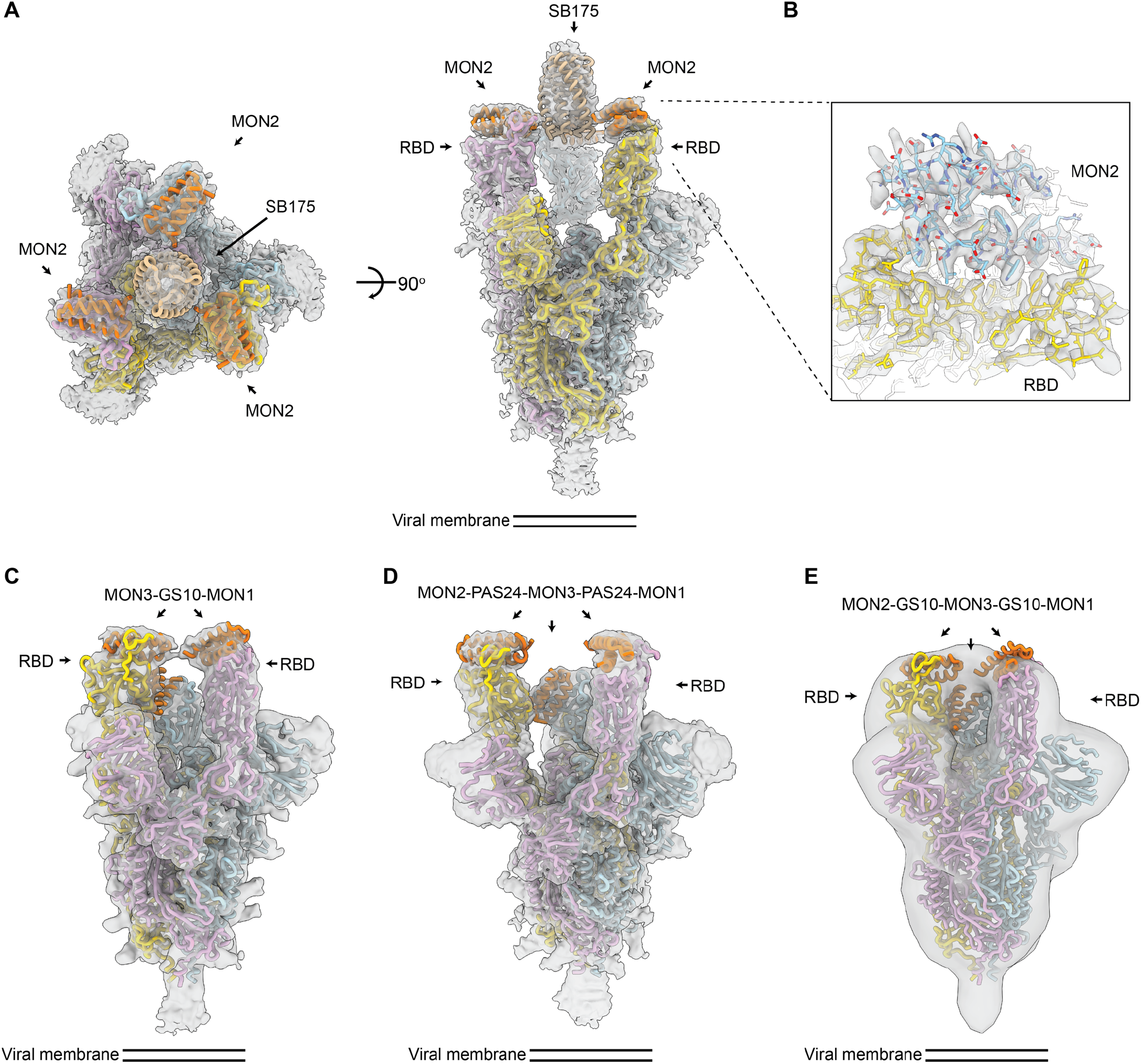
CryoEM structures of multivalent minibinders in complex with the SARS-CoV-2 S6P glycoprotein. (**A**) CryoEM map of TRI2-2 in complex with the S6P in two orthogonal orientations. (**B**) Zoomed-in view of the TRI2-2 and S6P complex interface obtained using local refinement of the RBD and TRI2-2. The RBD and MON2 built at 2.9 Å resolution are shown in yellow and blue, respectively. (**C**) CryoEM map of FUS31-G10 in complex with two RBDs. (**D**) CryoEM map of FUS231-P24 in complex with three RBDs. (**E**) Negative-stain EM map of FUS231-G10 in complex with S6P. The S trimer and minibinder models are placed in the whole map by rigid body fitting. EM density is shown as a transparent gray surface with a fitted atomic model. Spike protomers (PDB 7JZL) are shown in yellow, blue, and pink. Minibinders (PDB 7JZU, 7JZM, and MON2 atomic structure in this study) are shown in orange.

The cryoEM structures of the TRI2-2, FUS31-G10, and the FUS231-P24 constructs in complex with S6P were determined at resolutions of 2.8, 4.6, and 3.9 Å respectively (Fig. 2, Fig. S7-S10, Table S3). The TRI2-2/S6P cryoEM structure closely matched the TRI2-2 design, with all three RBDs in the open state bound to MON2 (Fig. 2A-B, Fig. S7). In the FUS31-G10 and S6P complex, FUS31-G10 is bound to two RBDs, both appearing to adopt the open conformation upon binding (Fig. 2C, Fig S9). The distance between the two RBDs in the open conformation is significantly closer in the FUS31-G10 structure than in the FUS231-P24 structure (Fig. 2C and D), suggesting that the bound minibinder pulls the RBDs closer together, in agreement with the shorter linkers used in the former minibinder construct. In the structure, two molecules of FUS31-G10 are bound to a single S trimer with the third RBD being occupied by a second FUS31-G10 molecule. In the structure of FUS231-P24 bound to S6P, FUS231-P24 bound to three RBDs. We were able to make tentative assignments of minibinder domain identities by rigid body fitting, with MON1 binding a closed conformation RBD and MON2 and MON3 binding to open conformation RBDs in a hypothetical structural model. Although the linkers are disordered in the cryoEM map, precluding definitive assessment of the connectivity between each minibinder module, the distances between the termini of the minibinder domains is compatible with the computational design models and strongly suggestive of engagement of either 2 (FUS31-G10) or 3 of the RBDs (FUS231-P24) in a single S trimer by the multivalent minibinders.

The structure of MON2 in complex with the RBD has not previously been determined. Starting from the TRI2-2/S6P cryoEM data, we improved the RBD/MON2 densities using focused classification and local refinement yielding maps at 2.9 Å resolution enabling visualization of the interactions formed by MON2 with the RBD. Superimposition of the design MON2 model to the corresponding cryoEM structures, using the RBD as reference, shows that the MON2 minibinder closely matched the design MON2 model with backbone Cɑ RMSD of 1.3 Å (Fig. S7E, and F). Together with previous structures of MON1 and MON3 (*19*), these data illustrate the accuracy with which both protein scaffolds and protein binding interfaces can now be computationally designed.

### Multivalent minibinder enables rapid detection of SARS-CoV-2 S

Having confirmed the binding mode of the FUS231 proteins via cryoEM, we sought to design an S trimer sensor, reasoning that the high affinity binding of the FUS231 proteins to the S trimer could make a useful diagnostic (*32*). We hypothesized that it would be possible to construct a bioluminescence resonance energy transfer (BRET) sensor for S trimer, where simultaneous engagement of all three minibinders in FUS231 with the S trimer would bring the N- and C-termini close enough together to enable efficient energy transfer. Towards this goal, we designed a BRET sensor based on FUS231-P12 with teLuc and mCyRFP3 fused to the N- and C-terminus of FUS231-P12 respectively (Fig. 3A) (*33, 34*). Upon binding of the sensor protein to a stabilized S trimer with 2 proline mutations (S2P) (*28, 32*), we observed a 350% increase in the 590 nm/470 nm BRET ratio, which was not observed when bound to the RBD alone, and determined the limit of detection to be 11 pM S2P (Fig. 3B, 3C, Fig. S10). The assay is easy to use, can be read out via mobile phone camera, and similar homogeneous assays have shown success in quantifying analytes in complex media like serum and saliva (*35*). Furthermore, these results support the proposed multivalent binding mode for the FUS231 proteins.

**Fig. 3.**
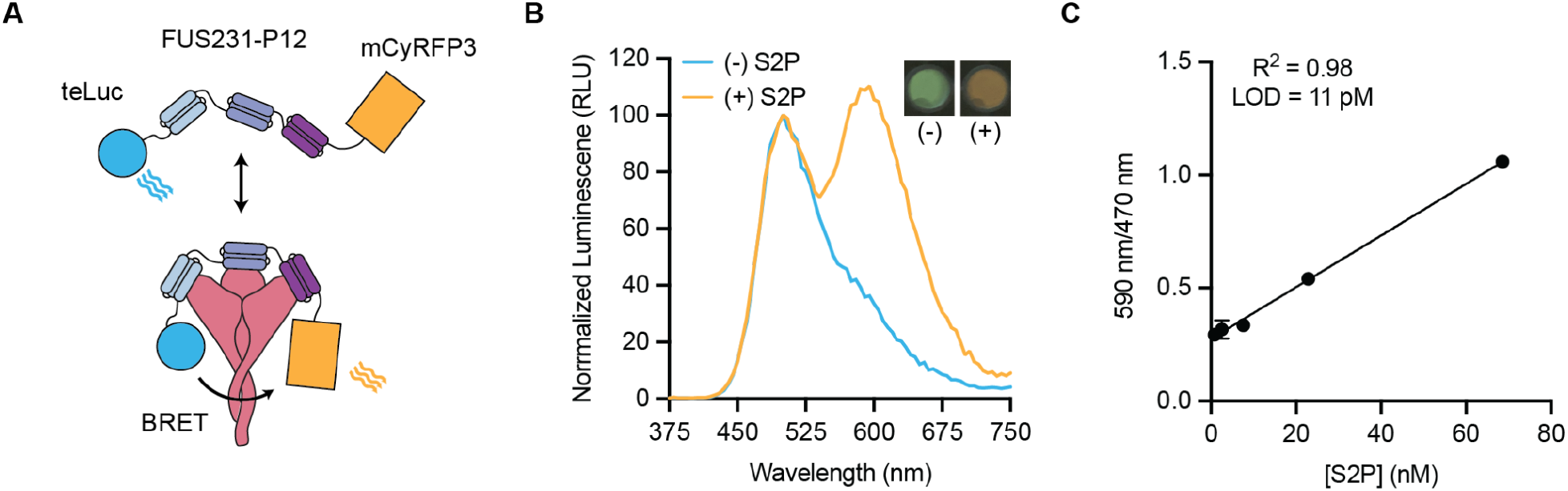
FUS231-P12 enables detection of SARS-CoV-2 S trimer via BRET. (**A**) Schematic representation of the BRET sensor, teluc-FUS231-P12-mCyRFP3, to detect S trimer. (**B**) Luminescence emission spectra and image of the BRET sensor (100 pM) in the presence (orange trace, 100 pM) and absence (blue trace) of S2P. Emission color change was observed using a mobile phone camera (inset top right). (**C**) Titration of S2P with 100 pM sensor protein (Mean ± SEM, n = 3 replicates from a single experiment).

### Multivalent minibinders bind tightly to circulating SARS-CoV-2 variants

We next evaluated the resiliency of the binding of multivalent minibinders to the previously identified MON1 and MON2 escape mutants as well as mutations present in the B.1.1.7, B.1.351, and P.1 SARS-CoV-2 variants. We first measured the off rate of the best multivalent minibinders using competition AlphaLISA with TRI2-1 against a panel of mutant spike proteins (Fig. 4A). Multivalent minibinders were fully bound to the mutant spike protein and subsequently were competed with 100-fold molar excess of untagged TRI2-1 to measure dissociation of the complex. The two-domain fusions (FUS23 and FUS31) showed little increased resilience to the tested point mutants. The three-domain fusions (FUS231) showed consistent binding to the tested mutants, indicating they are more resistant to mutation than their monomeric counterparts, though E406W, Y453R, and the combination of K417N, E484K, and N501Y mutations present in the B.1.351 S trimer increased the dissociation rate more than 100-fold. Consistent with these results, we also observed increased dissociation rates for the FUS231 proteins against the B.1.351 and P.1 spikes via Surface Plasmon Resonance (SPR) (Fig. S12). The TRI1 and TRI3 homotrimers showed similar mutational tolerance in the competition experiment, with the same E406W, Y453R, and B.1.351 mutations causing increased dissociation rates. Notably, the TRI2 proteins showed little dissociation after 24 hours against any of the tested S6P mutants.

**Fig. 4.**
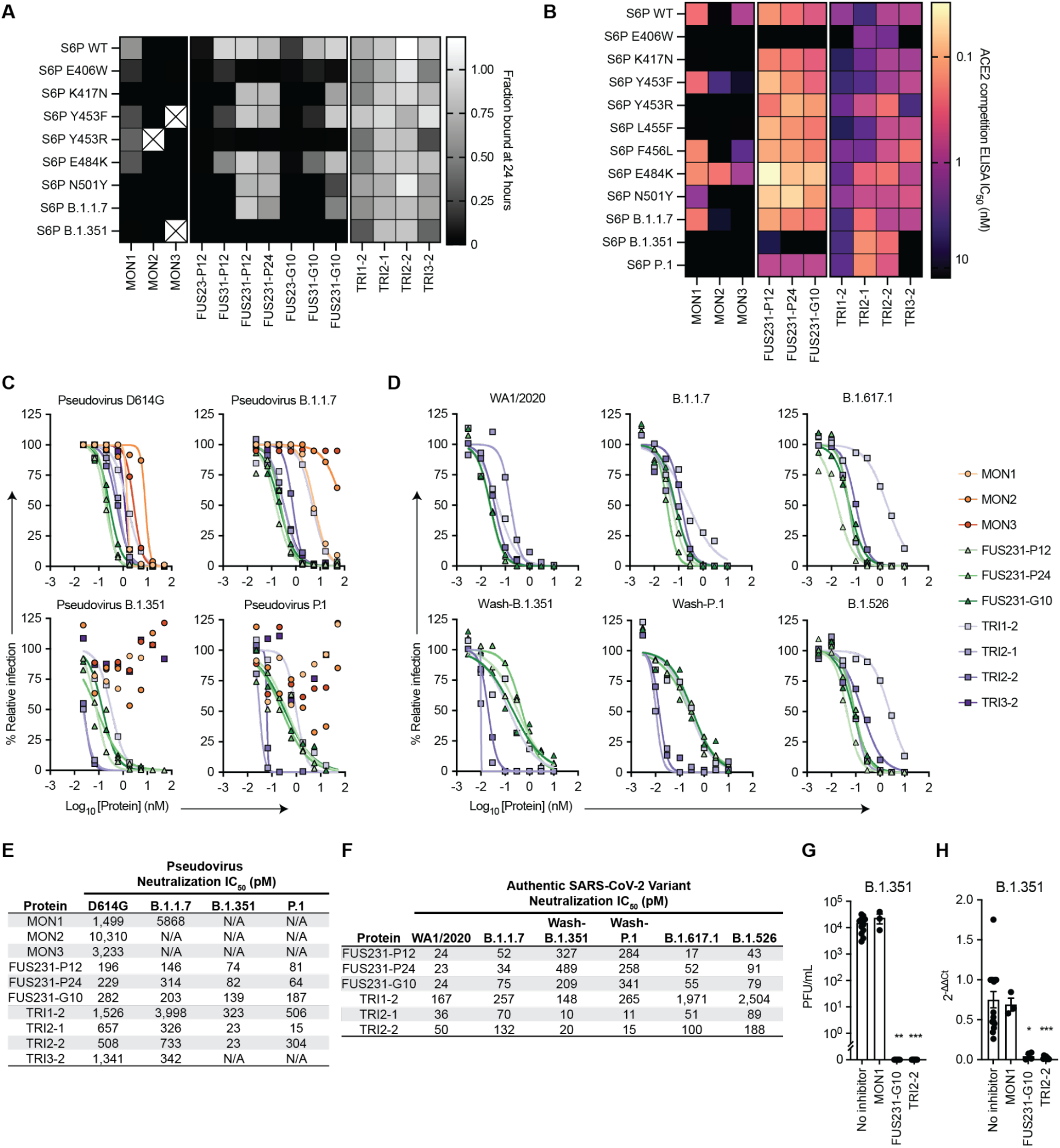
Multivalency enhances both the breadth and potency of neutralization against SARS-CoV-2 variants by minibinders. (**A**) Dissociation of minibinder constructs from S6P variants after 24 hours was measured via competition with untagged TRI2-1 using AlphaLISA (mean, n = 3 replicates from a single experiment). Cells containing an X indicate insufficient signal in the no competitor condition to quantify the fraction of protein bound. (**B**) Competition of minibinder constructs with ACE2 for S6P measured via ELISA (mean, n = 2). (**C**) Neutralization of SARS-CoV-2 pseudovirus variants by minibinder constructs (mean, n = 2). (**D**) Neutralization of authentic SARS-CoV-2 by minibinder constructs (mean, n = 2). (**E**) Table summarizing neutralization potencies of multivalent minibinder constructs against SARS-CoV-2 pseudovirus variants. N/A indicates an IC_50_ value above the tested concentration range and an IC_50_ greater than 50,000 pM. (**F**) Table summarizing neutralization potencies of multivalent minibinder constructs against authentic SARS-CoV-2 variants. (**G**) Neutralization of B.1.351 SARS-CoV-2 variant by minibinder constructs (0.3 µM) in human kidney organoids (n = 3 to 12: Kruskal-Wallis test: ** P < 0.01, *** P < 0.001). (**H**) Relative gene expression of SARS-CoV-2 envelope protein (SARS-CoV2-E) in kidney organoids post viral infection with and without multivalent mini binders (0.3 µM) (n = 3 to 15: Kruskal-Wallis test: * P < 0.05, *** P < 0.001).

Following binding measurements to assess dissociation rates, we screened the ability of the top multivalent minibinders to bind mutant S trimers by ACE2 competition ELISA, which correlates with neutralization potency (*36*). The minibinders were pre-incubated with the S6P variants before binding to immobilized ACE2 (Fig. 4B and Fig. S13). In line with the deep mutational scan, E406W, K417N, and Y453R, in addition to several other mutations, impaired binding. Two mutations, Y453F and E484K, improve MON2 binding, consistent with MON2 mimicry of the ACE2 interaction interface (*37*). Compared to the monovalent minibinders, we observed reduced effects of mutations in the competition IC_50_ values of the FUS231 and TRI2 minibinders and to a lesser extent of the TRI1 and TRI3 minibinders against the tested S6P variants except for E406W (Fig. 4B, Fig. S13D). Additionally, TRI2 minibinders show improved competition against N501Y containing spikes, consistent with improved ACE2 affinity to N501Y containing RBDs (*37, 38*).

### Multivalent minibinders potently neutralize circulating SARS-CoV-2 variants

To investigate the efficacy of the multivalent minibinders for preventing viral infection, we performed neutralization assays with the inhibitors using both a pseudotyped lentivirus and authentic SARS-CoV-2 variants. Against pseudoviruses displaying spike proteins corresponding to the B.1.1.7, B.1.351, and P.1 variants, all three monomer minibinders showed reduced neutralization capacity compared to the Wuhan-1 strain, whereas many of the multivalent minibinders were less affected in an ACE2 overexpressing cell line (Fig. 4C, E). The same proteins also were evaluated against pseudoviruses containing the E406W, L452R, and Y453F mutations which again had little impact on neutralization for most multivalent minibinders tested (Fig. S14). This suggests that the increase in affinity from multivalency improved neutralization breadth in a pseudovirus neutralization assay. The top neutralizing minibinders from this screen were selected for studies against authentic SARS-CoV-2 viruses including a historical WA1/2020 strain, B.1.1.7, B.1.526 (S477N), and B.1.617.1 natural isolates, and chimeric WA1/2020 strains encoding spike genes corresponding to those of B.1.351 (Wash-B.1.351), and P.1 (Wash-P.1) variants. Again, the top candidates maintained pM-range IC_50_ values (Fig. 4D and F). Notably, the TRI2 proteins maintained potent neutralization across all tested variants.

While Vero cells are useful for neutralization studies, they may not fully reflect the human antiviral response. Recent findings underscore the relevance of using non-transformed human organoid models for SARS-CoV-2 research (*39*). SARS-CoV-2 has been shown to infect and readily replicate in human kidney organoids, specifically targeting kidney tubular epithelial cells expressing ACE2 receptors, responsible for viral entry (*40*). Therefore, we generated kidney organoids from (H9) human embryonic stem cell line (*41*) (Fig. S15) and evaluated the ability of the multivalent minibinders to prevent SARS-CoV-2 viral entry in human tissues. Human kidney organoids were protected against the B.1.351 variant when the virus was pre-incubated with designed multivalent mini binders FUS231-G10 and TRI2-2, but not with MON1 (Fig. 4G). RT-qPCR analysis of RNA from the kidney organoids also showed reduced SARS-CoV-2 envelope protein (SARS-CoV2-E) gene expression after treatment with either FUS231-G10 or TRI2-2 (Fig. 4H). These data show that designed multivalent minibinders are potent neutralizers of the B.1.351 variant in a human organoid system that reflects the human antiviral response.

### Multivalent minibinders resist viral escape

Given the promising neutralization data, we subsequently tested the ability of the multivalent minibinders to resist viral escape mutations in the S trimer (Fig. 5A-B) (*2*). Plaque assays were performed with a VSV-SARS-CoV-2 S chimera on Vero E6 cells with minibinders included in the overlay to halt replication of non-resistant viruses. In positive control wells, treatment with a neutralizing antibody (2B04) in the overlay resulted in multiple escape mutants in each plate (*42*). In contrast, for both FUS231-P12 and TRI2-2, no escape mutants were isolated in 36 replicate wells for each protein (Fig. S16). These data indicate that both the FUS231-P12 and TRI2-2 proteins are substantially more difficult to escape than a typical RBD-targeted neutralizing mAb.

**Fig. 5.**
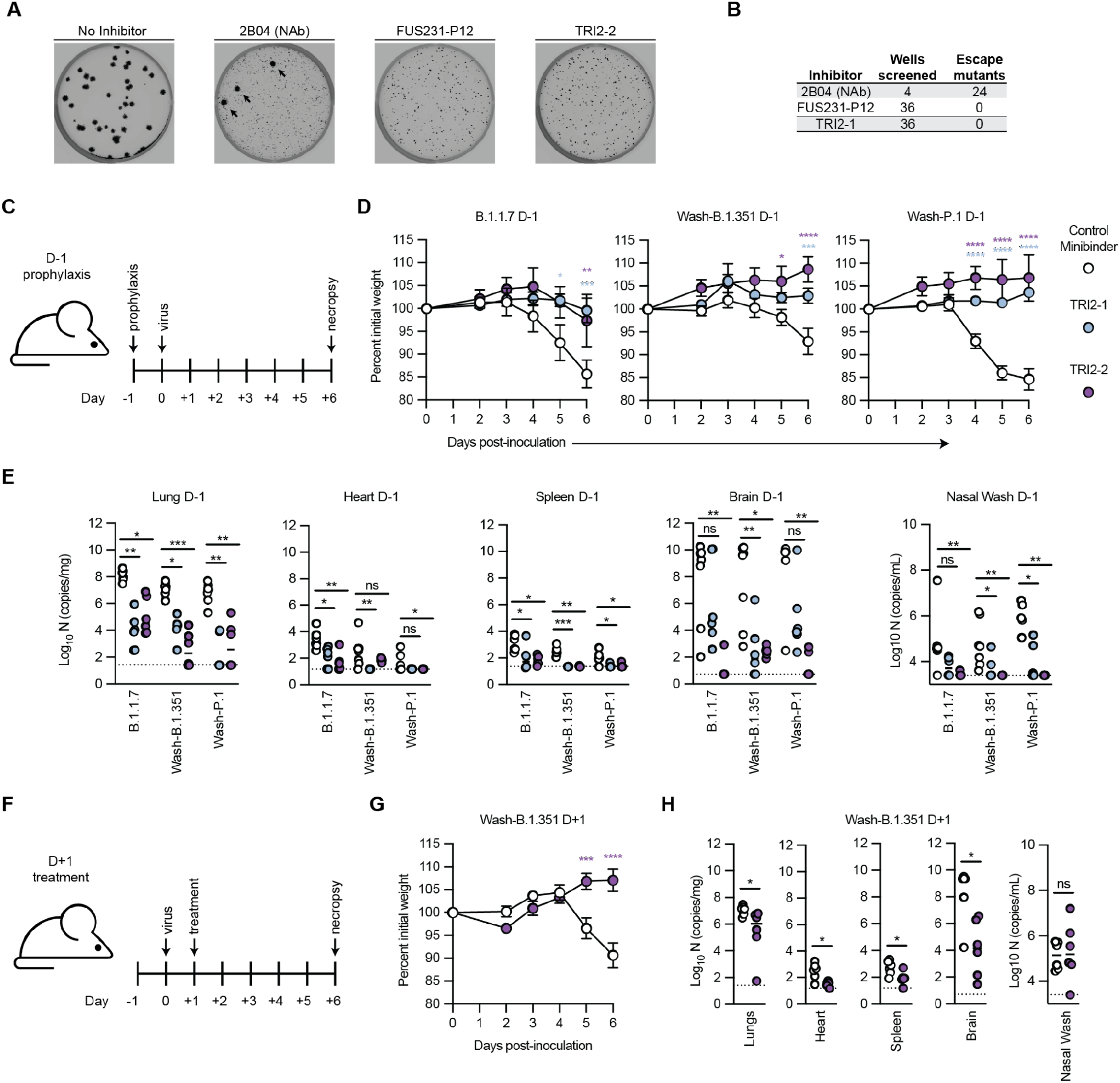
Top multivalent minibinder candidates are escape resistant and protect mice from SARS-CoV-2 infection via pre- and post-exposure intranasal administration. (**A**) Plaque assays were performed to isolate VSV-SARS-CoV-2 chimera virus escape mutants against a control neutralizing antibody (2B04) and the FUS231-P12 and TRI2-2 multivalent minibinders. Images are representative of 36 replicate wells per multivalent minibinder. Large plaques, highlighted by black arrows, are indicative of escape. (**B**) Table summarizing the results of the viral escape screen. (**C**-**E**) K18-hACE2-transgenic mice (n = 6/timepoint) were dosed with 50 μg of the indicated minibinder by i.n. administration (50 μl total) 24 h prior (D-1) to infection with 10^3^ focus forming units of SARS-CoV-2 variants B.1.1.7, Wash-B.1.351, or Wash-P.1 i.n. on Day 0. (**D**) Daily weight-change following inoculation (mean ± SEM; n = 6, two-way ANOVA with Sidak’s post-test: * P < 0.05, ** P < 0.01, *** P < 0.001, **** P < 0.0001). (**E**) At 6 days post infection (6 dpi) animals (n = 6/timepoint) were sacrificed and analyzed for the presence of SARS-CoV-2 viral RNA by RT-qPCR in the lung, heart, spleen, brain, or nasal wash (n = 6: Kruskal-Wallis test: ns, not significant, * P < 0.05, ** P < 0.01, *** P < 0.001). (**F**-**H**) K18-hACE2-transgenic mice (n = 6/timepoint) were dosed with 50 μg of the indicated minibinder by i.n. administration (50 μl total) 24 h after (D+1) infection with 10^3^ focus forming units of the SARS-CoV-2 Wash-B.1.351 variant on Day 0. (**G**) Daily weight-change following inoculation (mean ± SEM; n = 6, two-way ANOVA with Sidak’s post-test: * P < 0.05, ** P < 0.01, *** P < 0.001, **** P < 0.0001). (**H**) At 6 dpi, animals (n = 6/timepoint) were sacrificed and analyzed for the presence of SARS-CoV-2 viral RNA by RT-qPCR in the lung, heart, spleen, brain, or nasal wash (n = 6: Mann-Whitney test: ns, not significant, * P < 0.05, ** P < 0.01, *** P < 0.001).

### Multivalent minibinder confers protection in human ACE2-expressing transgenic mice

To determine the ability of our multivalent minibinders to prevent or treat SARS-CoV-2 infection *in vivo*, we evaluated pre-exposure prophylaxis or post-exposure therapy in highly susceptible K18-hACE2 transgenic mice (Fig. 5C-F). For prophylaxis, a single 50 μg dose (∼2.5 mg/kg) of TRI2-1 or TRI2-2 was administered via intranasal route one day prior to inoculation with 10^3^ focus forming units (FFU) of the indicated SARS-CoV-2 variant. In all cases, intranasal administration of TRI2-1 or TRI2-2 protected mice against SARS-CoV-2-induced weight loss. At 6 days post infection, viral burden in tissues was determined and shown to be reduced at almost all primary (lung and nasal wash) and secondary sites (heart, spleen, brain) of viral replication in TRI2-1 and TRI2-2 treated animals. To determine the therapeutic potential of our lead candidate, TRI2-2, we inoculated K18-hACE2 mice with 10^3^ FFU of Wash-B.1.351 and one day later, administered a single 50 μg dose of minibinder. Treatment with TRI2-2 protected against weight loss, and all tissues showed reduced viral burden except nasal washes. These results indicate that intranasal administration of TRI2-1 or TRI2-2 can protect as both pre-exposure prophylaxis and post-exposure therapy against SARS-CoV-2 infection in a stringent mouse model of disease.

## Conclusions

The coupling of structure-guided computational protein design, cell-free expression, and a competition-based off-rate screen enabled rapid identification and optimization of S trimer engaging multivalent inhibitors. These multivalent minibinders, whose designs were structurally validated by Cryo-EM, have several potential advantages over mAbs for the treatment of COVID-19. For example, they are amenable to large-scale and low-cost production in microorganisms like *E. coli,* they can readily be administered directly to the respiratory system, and their stability may obviate the need for cold chain storage. Our top candidate protein, TRI2-2, potently neutralizes historical and current SARS-CoV-2 variants of concern and provides prophylactic and therapeutic protection against all tested variants in highly susceptible K18-hACE2 transgenic mice. While multivalent RBD binders have been described previously (*43–52*), the trimeric ACE2 mimic TRI2-2 is the first miniprotein to engage the S protein trivalently with an inherently escape resistant binding interaction; the combination of trivalency and receptor mimicry (*36, 53, 54*) could be a useful approach for combating escape resistance. Looking forward, our integrated computational design and experimental screening pipeline for identifying geometrically matched arrays of receptor mimicking minibinders may provide a rapid and powerful strategy for developing protein-based medical countermeasures and diagnostic reagents against novel pathogens in the future.

## Acknowledgments

We thank Ashty S. Karim and Lauren Clark for their laboratory support during COVID-19 and Ali Ellebedy for providing antibodies.

## Author contributions

This was a community-based project that was only possible through unique combinations of expertise from multiple groups. As a result, there were many equal contributions and we would like to acknowledge that no single ordering of authors could have captured the importance of each contribution. A.C.H developed the cell-free DNA assembly and protein expression workflow, developed the off-rate multivalency screening assay, assisted in the optimization of the minibinders and selection of final candidates, manufactured the multivalent minibinders for screening and cell-based assays, screened the minibinders and multivalent minibinders in the off-rate assay and analyzed the results, and prepared the manuscript for submission. J.B.C. performed authentic virus neutralization assays and data analysis; assisted in minibinder prioritization and selection of final candidates; performed mouse *in vivo* experiments, tissue processing, virus quantification, and data analysis. Y.J.P. performed cryoEM experiments and analysis. L.Cao optimized the de novo minibinders and originated the multivalent project. L.Cao and A.C.H. originated the multi-domain fusions. L.Cao, L.S., and A.C.H. designed the multi-domain fusions. K.W. originated the homotrimers and designed and optimized the homotrimers. A.C.W. designed and piloted the minibinder-S6P-ACE2 competition assay and assisted with data analysis; organized reagents and prioritized and planned experiments; and performed pseudovirus neutralization assays and data analysis. Z.L. performed and analyzed the chimeric virus escape experiments. J.B. performed and analyzed the competition ELISA experiments, with the assistance of A.A.. H.W.Y, with the assistance of N.P., performed and analyzed the sensing experiments. S.S., L.H., Y.T.Z., and T-Y. H. performed and analyzed the kidney organoid experiments. T.N.S. performed and analyzed the RBD deep mutational scanning experiments, with the assistance of A.A.. I.G. and L.K. assisted in the optimization of minibinders. L.C. expressed multivalent minibinders for animal studies and expressed S6P variants. R.R. performed and analyzed heat purification experiments. L.B.G., W.L.M, and C.A.T, performed and analyzed BLI and SPR experiments. B.V. and A.K.G. assisted with expression and purification of minibinders and with the off-rate screening assay. L.A.V., R.E.C., B.Y., A.L.B., and N.M.K. assisted with authentic virus neutralization studies and mouse studies. S.B. designed SB175. A.L., N.E., G.U. assisted in the design of the homotrimers. A.L. assisted in the computational method development for homotrimers. N.E. and G.U. assisted in linker and oligomer selection. C.C. assisted with the expression of homotrimers. A.C.H., J.B.C., L.Cao, K.W., A.C.W., L.C., M.S.D., D.V, M.C.J, D.B. coordinated the research. M.G., B.F., B.R.L., J.D.B., H.R-B., S.P.J.W., L.S., M.S.D., D.V., M.C.J., D.B. supervised the research and secured funding. A.C.H., J.B.C., Y.J.P., L.Cao, A.C.W., K.W., J.E.B., M.S.D., R.R., S.B., L.S., H.W.Y., D.V., M.C.J., and D.B. wrote and edited the manuscript. All authors discussed the results and commented on the manuscript.

## Data and materials availability

Structural models and density maps have been deposited in the Protein Data Bank (PDB) and Electron Microscopy Data Bank (EMDB). Illumina sequencing data for the deep mutational scanning experiments are available on NCBI SRA, BioSample SAMN19925005. Data are available from the authors upon request.

## Supplementary Materials

Materials and Methods

Figs. S1 to S16

Tables S1 to S4

References (*55*–88)

