## Supplemental Materials for "Multivalent designed proteins protect against SARS-CoV-2 variants of concern"

**This PDF file includes:**

Materials and Methods
Figs. S1 to S16
Tables S1 to S3
Caption for Table S4

**Other Supplementary Materials for this manuscript include the following:**

Table S4 (.csv)

### Materials and Methods

Cell lines (Pseudovirus neutralization)

Expi293F cells (ThermoFisher) were grown in Expi293 Expression Medium (Gibco), cultured at 37°C with 8% $CO_2$ and shaking at 130 rpm. HEK293T/17 is a female human embryonic kidney cell line (ATCC). The HEK-ACE2 adherent cell line was obtained through BEI Resources, NIAID, NIH: NR-52511. All adherent cells were cultured at 37°C with 8% $CO_2$ in flasks with DMEM + 10% FBS (Hyclone) + 1% penicillin-streptomycin. Cell lines were not tested for mycoplasma contamination nor authenticated.

Cell lines (VSV SARS-CoV-2 Chimera escape selections)

Vero CCL81 and MA104 were obtained from ATCC and used as described previously (*55*).

Cell lines (SARS-CoV-2 neutralization)

Vero CCL81 (ATCC), Vero-TMPRSS2, and Vero-hACE2-TMPRSS2 (a gift of A. Creanga and B. Graham, NIH) were cultured at 37°C in Dulbecco's Modified Eagle medium (DMEM) supplemented with 10% fetal bovine serum (FBS), 10 mM HEPES pH 7.3, 1 mM sodium pyruvate, 1× non-essential amino acids, and 100 U/ml of penicillin–streptomycin. Additionally, Vero-TMPRSS2 and Vero-hACE2-TMPRSS2 cells were cultured in the presence of 5 µg/mL of blasticidin or puromycin, respectively. The WA1/2020 (2019n-CoV/USA_WA1/2020) isolate of SARS-CoV-2 was obtained from the US Centers for Disease Control (CDC). WA1/2020 stocks were propagated on Vero CCL81 cells and used at passage 6 and 7. The B.1.1.7, Wash-B.1.351, and Wash-B.1.1.28 viruses have been described previously (*7*, *56*). B.1.526 (S477N) and B.1.617.1 viruses were isolated from infected individuals. For all strains, infectious stocks were propagated by inoculating Vero CCL81 or Vero-TMPRSS2 cells. Supernatant was collected, aliquoted, and stored at -80°C. All work with infectious SARS-CoV-2 was performed in Institutional Biosafety Committee-approved BSL3 and A-BSL3 facilities at Washington University School of Medicine using positive pressure air respirators and protective equipment. All virus stocks were deep-sequenced after RNA extraction to confirm the presence of the anticipated substitutions.

LCB1 (MON1) and LCB3 (MON3) optimization

Site saturation mutagenesis data were collected for LCB1 and LCB3 using the method described previously (*19*). Beneficial mutations that showed increased binding with RBD were selected and used to construct a Resfile (*57*). 10,000 sequence design trajectories were performed using the Rosetta FastDesign protocol, constrained by the Resfile. Eight sequences were selected based on rosetta score and binding ddG for LCB1 and LCB3 respectively, while keeping the sequence diversity. Genes encoding the selected sequences were cloned into modified pET-29b(+) E. coli plasmid expression vectors (GenScript, N-terminal 8 His-tag followed by a TEV cleavage site) for checking the expression yield (the expression yield is often correlated with protein solubility and stability) and the binding with RBD was characterized with the AlphaLISA assay (see below), the sequences with high expression yield, as well as tight binding affinity with RBD were selected for downstream multivalent constructs design, with the names as LCB1_v2.2 and LCB3_v2.2.

Multivalent fusion constructs design

The CryoEM structures of LCB1 and LCB3 in complex with the spike protein, and the design model of AHB2 were used to determine the order of the monomers in the multivalent fusion constructs. The structures/model were firstly aligned using the spike protein as the reference and the order was determined with the criterion that a shorter linker had to be short when the two linked binders bound with the same spike simultaneously. Four different combinations, AHB2-LCB1, LCB3-LCB1, AHB2-LCB3, LCB3-AHB2, LCB3-AHB2-LCB1, and AHB2-LCB2-LCB1, were tested with either a flexible glycine-serine (GS) linker or a proline-alanine-serine (PAS) linker (*58*). The linker length was optimized using the AlphaLISA assay (see below).

Self-assembling homotrimer design: scaffold selection and backbone generation

De novo designed C3 symmetric protein scaffolds and de novo designed helical repeat proteins (DHRs) are the basis for the generation of trimeric mini-binders. 34 designed C3 scaffolds and 62 DHRs were selected after preliminary screening based on the geometry matching with the trimeric SARS-CoV-2 S protein and scaffold quality. Fusing the three components, i.e., C3 scaffolds, DHRs and designed mini-binders together through a modified version of the WORMS (*59*) software, 1096 trimeric binder backbones were generated. This was followed by steric clashing filtering (Rosetta centroid energy < 10), visual inspection in a molecular graphics viewer (Pymol), and sequence design for residues within 9 Å from the fused junction. 56 designs were selected for further validation. To ensure the matching geometry between the fused constructs and the SARS-CoV-2 S trimer with three RBD in the open state (PDB: 7CAK), the WORMS package was customized with the "stack" orientation implemented, where the two cyclic symmetrical axes from both C3 scaffolds and Spike trimer are aligned along Z axis.

Self-assembling homotrimer design: junction modification and flexibility insertion

Considering the strict geometric requirement, the dynamic nature of the S trimer and the low expression successful rate, two new approaches have been pursued to simplify the constructs and enhance the flexibility.

The first approach was creating semi-flexible binders. Starting from the high quality rigid fusion models with a relatively small size (<250 amino acids in total), DHR junctions were omitted. Instead, the last one or two helices from the C3 scaffolds were modified, truncated, or extended with one or two helices using blueprint-based backbone generation (*60*, *61*). The newly generated last helix was required to be within 12 Å to the first helix from mini-binders in models. (Glycine-Serine)x1 to (Glycine-Serine)x6 were modeled in Pymol as the flexible linker between the modified C3 scaffolds and mini-binders. Based on the success of this approach, the second was introduced to simplify the constructs further with short flexible linkers only. Four de novo C3 scaffolds and six native C3 scaffolds were selected as final candidates with the right geometry, some of which confirmed by the solved cryo-electron microscopy structures. The modification on the last helix was allowed within two to ten residues to be as geometrically compatible as previously. The sequence was optimized based on Rosetta combinatorial sequence optimization packages (*62–64*) and homology models.

Deep mutational scanning profiles of minibinder escape

Minibinder escape mapping experiments were performed in biological duplicate using a deep mutational scanning approach as previously described (*3*, *65*). Briefly, yeast-surface display libraries expressing 3,804 of the 3,819 possible amino acid mutations in the SARS-CoV-2 RBD (Wuhan-Hu-1 sequence, Genbank MN908947, residues N331-T531) were constructed in (*37*), and sorted to purge mutations that abolish RBD folding or ACE2 binding. Libraries were induced for

RBD surface expression and labeled with minibinder at the concentration of the EC90 for binding to yeast-displayed RBD ("sensitive" selection) or 400 ng/mL ("stringent" selection), the concentration used for prior selections of clinical antibodies (*3*, *66*), followed by secondary labeling with 1:100 FITC-conjugated anti-Myc (Immunology Consultants Lab, CMYC-45F), and 1:100 iFluor-647-conjugated mouse anti-His tag antibody (Genscript A01802). Library variants that escape miniprotein binding were sorted on a BD FACSAria II based on gates drawn from control populations labeled at 0.1x ("sensitive" selection) or 0.01x ("stringent" selection) the selection concentration, as shown in Fig. S2A-C. For each sample, >10 million $RBD^+$ cells were processed, minibinder-escape cells were sorted and grown overnight, plasmid purified, and sequenced on a HiSeq 2500. Escape fractions were computed from sequencing counts exactly as described in Starr et al. (*3*). Illumina sequencing counts are available from the NCBI SRA (BioProject SAMN19925005). All code and analysis steps are described on GitHub: https://github.com/jbloomlab/SARS-CoV-2-RBD_MAP_minibinders. A table reporting all mutation escape fractions is available: https://github.com/jbloomlab/SARS-CoV-2-RBD_MAP_minibinders/blob/main/results/supp_data/IPD_ligands_raw_data.csv.

Cell-free DNA assembly and CFPS template preparation

Proteins to be manufactured via CFPS were codon optimized using the IDT codon optimization tool and ordered as gblocks containing the pJL1 5' (tttgtttaactttaagaaggagatatacat) and 3' (gtcgaccggctgctaacaaagcccgaaagg) Gibson assembly overhangs. DNA was resuspended at a concentration of 50 ng/μL.

Linearized pJL1 plasmid backbone (Addgene plasmid # 69496) was ordered as a gblock from IDT (see below) and amplified using the pJL1_F (gtcgaccggctgcta) and pJL1_R (atgtatatctccttcttaaagttaaacaaaattatttcta) primers via PCR using Q5 Hot Start DNA polymerase (NEB, M0493L) following manufacturer instructions. Amplified pJL1 backbone was purified using the DNA Clean and Concentrate Kit (Zymo Research, D4006) and diluted to a concentration of 50 ng/μL.

Linearized pJL1 plasmid backbone:
gtcgaccggctgctaacaaagcccgaaaggaagctgagttggctgctgccaccgctgagcaataactagcataaccccttggggcctcta
aacgggtcttgaggggttttttgctgaaagccaattctgattagaaaaactcatcgagcatcaaatgaaactgcaatttattcatatcaggattat
caataccatatttttgaaaaagccgtttctgtaatgaaggagaaaactcaccgaggcagttccataggatggcaagatcctggtatcggtctgc
gattccgactcgtccaacatcaatacaacctattaatttcccctcgtcaaaaataaggttatcaagtgagaaatcaccatgagtgacgactgaat
ccggtgagaatggcaaaagcttatgcatttctttccagacttgttcaacaggccagccattacgctcgtcatcaaaatcactcgcatcaaccaa
accgttattcattcgtgattgcgcctgagcgagacgaaatacgcgatcgctgttaaaaggacaattacaaacaggaatcgaatgcaaccggc
gcaggaacactgccagcgcatcaacaatattttcacctgaatcaggatattcttctaatacctggaatgctgttttcccggggatcgcagtggt
gagtaaccatgcatcatcaggagtacggataaaatgcttgatggtcggaagaggcataaattccgtcagccagtttagtctgaccatctcatct
gtaacatcattggcaacgctacctttgccatgtttcagaaacaactctggcgcatcgggcttcccatacaatcgatagattgtcgcacctgattg
cccgacattatcgcgagcccatttatacccatataaatcagcatccatgttggaatttaatcgcggcttcgagcaagacgtttcccgttgaatat
ggctcataacaccccttgtattactgtttatgtaagcagacagttttattgttcatgatgatatatttttatcttgtgcaatgtaacatcagagattttga
gacacaacgtgagatcaaaggatcttcttgagatcctttttttctgcgcgtaatctgctgcttgcaaacaaaaaaaccaccgctaccagcggtg
gtttgtttgccggatcaagagctaccaactctttttccgaaggtaactggcttcagcagagcgcagataccaaatactgttcttctagtgtagcc
gtagttaggccaccacttcaagaactctgtagcaccgcctacatacctcgctctgctaatcctgttaccagtggctgctgccagtggcgataa
gtcgtgtcttaccgggttggactcaagacgatagttaccggataaggcgcagcggtcgggctgaacggggggttcgtgcacacagcccag
cttggagcgaacgacctacaccgaactgagatacctacagcgtgagctatgagaaagcgccacgcttcccgaagggagaaaggcggac

aggtatccggtaagcggcagggtcggaacaggagagcgcacgagggagcttccagggggaaacgcctggtatctttatagtcctgtcgg gtttcgccacctctgacttgagcgtcgatttttgtgatgctcgtcaggggggcggagcctatggaaaaacgccagcaacgcgatcccgcga aattaatacgactcactatagggagaccacaacggtttccctctagaaataattttgtttaactttaagaaggagatatacat

Gibson assembly was used to assemble protein open reading frame DNA with the pJL1 backbone following the published protocol with the addition 3.125 μg/mL of ET SSB (NEB, product no. M2401S) (*67*, *68*). 20 ng of purified, linearized pJL1 backbone and 20 ng of the protein open reading frame insert were combined in 2 μL Gibson assembly reactions and incubated at 50°C for 30 minutes. Similar to other published methods (*21*, *69*), the unpurified assembly reactions were diluted in 40 μL of nuclease free water (Fisher Scientific, AM9937) and 1 μL of the diluted reaction were used as the template for a PCR to generate linear expression templates (LETs) for CFPS. Linear expression templates were amplified via PCR using the pJL1_LET_F (ctgagatacctacagcgtgagc) and pJL1_LET_R (cgtcactcatggtgatttctcacttg) primers and the Q5 Hot Start DNA polymerase (NEB, M0493L) following manufacturer instructions.

<u>CFPS cell extract preparation</u>
Cell extracts for CFPS reactions were prepared from BL21 Star$^{TM}$ (DE3) as previously described (*70*).

<u>CFPS reactions and purification</u>
CFPS reactions were assembled as previously described except for the DNA template (*70*). Unpurified linear expression templates in PCR reaction buffer were added to the CFPS reaction at 6.66% v/v to drive protein synthesis. CFPS reactions were run in well plates at 30°C for 14-20 h. 100-200 μL volume reactions, to produce protein for AlphaLISA off-rate screening, were run in 24-well plates (Falcon, 351147) and purified via StrepTactinXT spin columns (IBA, 2-4151-000) following manufacturer instructions. Post purification, proteins were extensively dialyzed (Fisher Scientific, 69552) into 50 mM HEPES pH 7.4, 150 mM NaCl.

2 mL reactions, to produce protein for ELISA, neutralization, and VSV escape studies, were run in 6-well plates (Costar, 3736) and purified via gravity flow using StrepTactinXT Superflow high capacity resin (IBA, 2-4030-010). CFPS reactions were incubated with resin on an end-over-end rotator for 30 min at room temperature. The resin was spun down at 2,500 x *g* for 2 min and the supernatant was removed. Resin was resuspended in Buffer W (IBA, 2-1003-100) and loaded onto a gravity flow column (BioRad, 7321010). Resin was washed with 20X resin volumes of Buffer W and proteins were eluted via the addition of Buffer BXT (IBA, 2-1042-025). Following purification, EDTA and CHAPS were added to a concentration of 10 mM and 4 mg/mL, respectively to aid in the removal of endotoxin by dialysis (*71*). Proteins were dialyzed (Fisher Scientific, 87724) into 1L of endotoxin-free 1x PBS (Fisher Scientific, SH30256LS) in a 4 L glass beaker. Glass beakers were baked at 240°C for 12 h to degrade endotoxin prior to use. Samples were dialyzed at 4°C for > 6 h. At least two additional rounds of dialysis with the addition of EDTA and CHAPS to concentrations of 10 mM and 4 mg/mL, respectively, were performed to remove endotoxin. Endotoxin was quantified using the Pierce Chromogenic Quant Kit (Fisher Scientific, A39553) and samples with less than 10 EU/mL were used for cell-based assays.

<u>Expression and purification of competitor proteins for AlphaLISA experiments</u>
Proteins used as competitors in the AlphaLISA off-rate screen were cloned into pET28a,

transformed into BL21 Star$^{TM}$ DE3, plated on LB agar, and cultured overnight at 37°C. 1 L of Overnight Express TB (Fisher Scientific, 71491-4) was inoculated by scraping all colonies on a transformation plate and cultured at 37°C in 2.5 L tunair flasks (IBI Scientific, SS-8003) at 220 rpm overnight. Cells were harvested, resuspended at a ratio of 1 g cell mass to 4 mL resuspension buffer (50 mM HEPES pH 7.5, 500 mM NaCl, 1X HALT protease inhibitor without EDTA (Fisher Scientific, 78429), 1 mg/mL lysozyme, 62.5 U/mL cell suspension of benzonase (Sigma Aldrich, E1014-25KU)) and lysed using an Avestin B15 homogenizer at 21,000 PSI. Lysate was spun down 14,000 x *g* for 10 min and the clarified supernatant was incubated with Ni-NTA Agarose (Qiagen, 30230) for 60 min on an end-over-end shaker. Resin was spun down 2,500 x *g* for 2 min, supernatant removed, resuspended in wash buffer (50 mM HEPES pH 7.5, 500 mM NaCl, 50 mM Imidazole), loaded on a gravity flow column, and subsequently washed with 20X resin volumes of wash buffer. Protein was eluted using elution buffer (50 mM HEPES pH 7.5, 500 mM NaCl, 500 mM Imidazole) and exchanged into 50 mM HEPES pH 7.4, 150 mM NaCl using PD-10 desalting columns (Cytvia, 17-0851-01).

His tags were removed via cleavage by ProTEV Plus (Promega, V6102). Prior to cleavage, 10% v/v glycerol was added to the protein. ProTEV Plus was added to a concentration of 0.5 U/µg purified protein and DTT was added to a concentration of 1 mM. Cleavage reactions were carried out at 30°C for 4 h. Free his tag and ProTEV Plus were removed by incubating with Ni-NTA Agarose for 1 h at 4°C and collecting the supernatant. Proteins were subsequently concentrated to > 1mg/mL (Millipore, UFC800396). His tag removal was validated via SDS PAGE and the AlphaScreen Histidine (Nickel Chelate) Detection Kit (Perkin Elmer, 6760619C).

<u>Expression and purification of SARS-CoV-2 S proteins</u>
S2P (*28*, *32*), S6P (*23*), and S6P variants were produced in Expi293F cells grown in suspension using Expi293F expression medium (Life Technologies) at 33°C, 70% humidity, 8% CO2 rotating at 150 rpm. The cultures were transfected using PEI-MAX (Polyscience) with cells grown to a density of 3.0 million cells per mL and cultivated for 3 days. Supernatants were clarified by centrifugation (5 min at 4000 rcf), addition of PDADMAC solution to a final concentration of 0.0375% (Sigma Aldrich, #409014), and a second spin (5 min at 4000 rcf).

Proteins were purified from clarified supernatants via a batch bind method where each clarified supernatant was supplemented with 1 M Tris-HCl pH 8.0 to a final concentration of 45 mM and 5 M NaCl to a final concentration of ~310 mM. Talon cobalt affinity resin (Takara) was added to the treated supernatants and allowed to incubate for 15 min with gentle shaking. Resin was collected using vacuum filtration with a 0.2 µm filter and transferred to a gravity column. The resin was washed with 20 mM Tris pH 8.0, 300 mM NaCl, and the protein was eluted with 3 column volumes of 20 mM Tris pH 8.0, 300 mM NaCl, 300 mM imidazole. The batch bind process was then repeated and the first and second elutions combined. SDS-PAGE was used to assess purity. IMAC elutions were concentrated to ~1 mg/mL and dialyzed three times into 50 mM Tris pH 8, 150 mM NaCl, 0.25% L-Histidine in a hydrated 10K molecular weight cutoff dialysis cassette (Thermo Scientific). Due to inherent instability, S2P was immediately flash frozen and stored at −80°C.

<u>Competition-based off-rate screening via AlphaLISA</u>
AlphaLISA reactions were carried out in 50 mM HEPES pH 7.4, 150 mM NaCl, 1 mg/mL BSA,

and 0.015% v/v TritonX-100 (hereafter referred to as Alpha buffer). All components were dispensed using an Echo 525 liquid handler from a 384-Well Polypropylene 2.0 Plus microplate (Labcyte, PPL-0200) using the 384PP_Plus_GPSA fluid type. All AlphaLISA reactions were performed in a ProxiPlate-384 Plus (Perkin Elmer, 6008280). AlphaLISA StrepTactin donor beads, to capture StrepII or TwinStrep-tagged minibinder variants, (Perkin Elmer, AS106) and AlphaLISA Anti-6x-his, to capture 6xhis-tagged S6P or RBD, (Perkin Elmer, AL178C) were combined to prepare a 4X stock in Alpha buffer immediately prior to use and added to the proteins to yield a concentration of 0.08 mg/mL donor beads and 0.02 mg/mL acceptor beads in the final reaction.

Multivalency screening experiments were carried out at a final concentration of 2.5 nM S6P, 2.5 nM minibinder variant, and 250 nM of the specified untagged competitor in Alpha buffer. First, the minibinder variant and S6P were diluted to 1.33x final concentration (adjusted for the later addition of competitor) in 120 µL in a 384-Well Polypropylene 2.0 Plus plate and sealed (BioRad, MSB1001). Samples were allowed to fully associate for 12-16 h at 20°C. Next, conditions were split in half and the same volume of either buffer or 250 nM (100x molar excess) untagged competitor were added to achieve 1.33x final concentration of all components (competitor was previously concentrated to achieve less than 5% volume change at this step). Samples were then incubated for the specified time, with replicates measured by dispensing 1.5 µL of each condition and 0.5 µL of 4X Alpha bead stock via the Echo 525 liquid handler. Plates were immediately spun down following the dispense and sealed (BioRad, MSB1001). Reactions were incubated with beads for 1 h for 2 h dissociation time points and up to 2 h for longer dissociation time points before measurement (bead incubation time was included in the specified timepoints). AlphaLISA measurements were taken on a Tecan Infinite M1000 Pro using the AlphaLISA filter with an excitation time of 100 ms, an integration time of 300 ms and a settle time 20 ms. Prior to measurement, plates were allowed to equilibrate inside the instrument for 10 min. Fraction of protein bound was determined by subtracting the average background bead signal and then dividing the plus competitor by the minus competitor condition. Values below zero after background subtraction were set to zero. Conditions with signal in the no competitor condition within 3 standard deviations of the background were not analyzed. Prism 9 (GraphPad) was used to plot the data.

Monovalent optimization experiments were performed in the same manner as multivalency screening experiments. S6P and RBD comparison experiments were carried out in the same manner as the multivalency screening experiments except for concentrations of 5 nM of S6P or RBD (Sino Biological, 40592-V08H), 5 nM minibinder variant, and 500 nM (100x molar excess) of the specified untagged competitor in the final reactions. In S6P variant experiments, reactions were carried out in the same manner as the multivalency screening experiments.

<u>Size exclusion chromatography with multi-angle light scattering (SEC MALS)</u>

Using an Agilent 1200 HPLC, we ran > 100 µg of the indicated sample in 50mM Tris-HCL, 150mM NaCl pH 8.0 at 0.35mL/min over an Agilent Bio-SEC 3 300A column in line with a Heleos multi-angle static light scattering and an Optilab T-rEX detector (Wyatt Technology Corporation). The data was then analyzed using ASTRA (Wyatt Technologies) to calculate the weighted average molar mass (Mw) of the selected species and the number average molar mass (Mn) to determine monodispersity by polydispersity index (PDI) = Mw/Mn.

Negative stain electron microscopy

The SARS-CoV-2 HexaPro Spike protein (S6P) was produced in HEK293F cells grown in suspension using FreeStyle 293 expression medium (Life technologies) at 37˚C in a humidified 8% CO2 incubator rotating at 130 rpm. The cultures were transfected using PEI (9 μg/mL) with cells grown to a density of 2.5 million cells per mL and cultivated for 3 d. The supernatants were harvested and cells resuspended for another 3 d, yielding two harvests. Spike proteins were purified from clarified supernatants using a 5 mL Cobalt affinity column (Cytiva, HiTrap TALON crude), concentrated and flash frozen in a buffer containing 20 mM Tris pH 8.0 and 150 mM NaCl prior to analysis.

10 μM SARS-CoV-2 Spike was incubated with 13 μM minibinders for 1 h at room temperature. Samples were diluted to 0.01 mg/mL immediately prior to adsorption to glow-discharged carbon-coated copper grids for ~30 sec prior to a 2% uranyl formate staining. Micrographs were recorded using the Leginon on a 120 KV FEI Tecnai G2 Spirit with a Gatan Ultrascan 4000 4k x 4k CCD camera at 67,000 nominal magnification. The defocus ranged from -1.0 to -2.0 μm and the pixel size was 1.6 Å.

Cryo-electron microscopy

SARS-CoV-2 HexaPro S (S6P) at 1.2 mg/mL was incubated with 1.2 fold molar excess of recombinantly purified TRI2-2, FUS31-G10, or FUS231-P24 at 4°C before application onto a freshly glow discharged 2.0/2.0 UltrAuFoil grid (200 mesh). Plunge freezing used a vitrobot MarkIV (ThermoFisher Scientific) using a blot force of 0 and 6.5 second blot time at 100% humidity and 23°C.

For the S6P/TRI2-2 data set, Data were acquired using an FEI Titan Krios transmission electron microscope operated at 300 kV and equipped with a Gatan K3 direct detector and Gatan Quantum GIF energy filter, operated in zero-loss mode with a slit width of 20 eV. Automated data collection was carried out using Leginon at a nominal magnification of 105,000x with a pixel size of 0.4215 Å. The dose rate was adjusted to 15 counts/pixel/s, and each movie was acquired in super-resolution mode fractionated in 75 frames of 40 ms. 5,991 micrographs were collected with a defocus range between -0.5 and -2.5 μm. Movie frame alignment, estimation of the microscope contrast-transfer function parameters, particle picking, and extraction were carried out using Warp (*72*) (Fig S7 and S8).

For the S6P/FUS31-G10 data set, data were acquired on an FEI Titan Krios transmission electron microscope operated at 300 kV equipped with a Gatan K2 Summit direct detector and Gatan Quantum GIF energy filter, operated in zero-loss mode with a slit width of 20 eV. Automated data collection was carried out using Leginon (*73*) at a nominal magnification of 130,000x with a pixel size of 0.525 Å. The dose rate was adjusted to 8 counts/pixel/s, and each movie was acquired in counting mode fractionated in 50 frames of 200 ms. 1000 micrographs were collected in a single session with a defocus range between -0.5 and -2.5 μm (Fig S9).

For the S6P/FUS231-P24 data set, data were acquired on an FEI Glacios transmission electron microscope operated at 200 kV equipped with a Gatan K2 Summit direct detector. Automated data collection was carried out using Leginon at a nominal magnification of 36,000x with a pixel size of 1.16 Å. The dose rate was adjusted to 8 counts/pixel/s, and each movie was acquired in counting mode fractionated in 50 frames of 200 ms. 1,663 micrographs were collected in a single session with a defocus range between -0.5 and -2.5 μm (Fig S10).

For the S6P/TRI2-2, S6P/FUS31-G10, and S6P/FUS231-P24 datasets, two rounds of reference-free 2D classification were performed using CryoSPARC (*74*) to select well-defined particle images. These selected particles were subjected to two rounds of 3D classification with 50 iterations each (angular sampling 7.5˚ for 25 iterations and 1.8˚ with local search for 25 iterations), using our previously reported closed SARS-CoV-2 S structure as initial model (PDB 6VXX) in Relion (*75*). 3D refinements were carried out using non-uniform refinement along with per-particle defocus refinement in CryoSPARC. Selected particle images were subjected to the Bayesian polishing procedure implemented in Relion3.0 before performing another round of non-uniform refinement in CryoSPARC followed by per-particle defocus refinement and again non-uniform refinement (*76*). To further improve the density of the TRI2-2, the particles were then subjected to focus 3D classification without refining angles and shifts using a soft mask on Receptor binding domain (RBD) and TRI2-2 region with a tau value of 60 in Relion. Particles belonging to classes with the best resolved local density were selected and subject to local refinement using CryoSPARC. Local resolution estimation, filtering, and sharpening were carried out using CryoSPARC. Reported resolutions are based on the gold-standard Fourier shell correlation (FSC) of 0.143 criterion and Fourier shell correlation curves were corrected for the effects of soft masking by high-resolution noise substitution. UCSF Chimera (*77*) and Coot (*78*) were used to fit atomic models into the cryoEM maps. Spike-RBD/TRI2-2 model was refined and relaxed using Rosetta (*79*) using sharpened and unsharpened maps (Table S2, Fig S8, S9, and S10).

<u>BRET sensor for SARS-CoV-2 S detection</u>

Synergy Neo2 plate reader (Biotek) was used for all luminescent assays. 10 μL of 1 nM teluc-FUS231-P12-mCyRFP3, 10 μL of serial diluted 10X spike protein (final concentrations range from 1.85 nM to 0.85 pM), and 30 μL of buffer (25 mM Tris, pH8, 50mM NaCl) were mixed and incubated for 10 min at room temperature. Diphenylterazine stock solution was prepared as previously described (*33*). Then, 50 μL of diluted diphenylterazine solution (60 μM) was added to each well. The luminescence spectra were collected under the monochromator mode with 0.1 s integration and 5 nm increments from 400 to 750 nm. The luminescent image was taken by an iPhone8 camera. To record the emission ratio, luminescence signals were acquired by a filter mode with a two channel cube (470/40 nm and 590/35 nm). The emission ratios were calculated from 590 nm/470 nm channels directly. The linear region of ratiometric responses was extracted and a linear regression curve was plotted, which was used to derive the standard deviation (s.d.) of the response and the slope of the calibration curve (S). The limit of detection was determined as 3 s.d. above background signal.

<u>Minibinder/RBD interaction kinetics via bio-layer interferometry (BLI)</u>

The Octet® HTX instrument was used to determine affinities of minibinders to RBD-HuFc (Acro Bio, #SPD-C5255) and monovalent soluble RBD (sol-RBD) (obtained from the Institute of Protein Design at University of Washington). Prior to RBD-HuFc affinity measurements, Anti-Human Fc Capture (AHC) biosensors (Sartorius, #18-5064) were pre-hydrated for 10 min in Octet running buffer containing 10 mM Tris (Fisher, #BP152-1), 150 mM NaCl (Fisher, #S271-10), 1 mM CaCl2 (Fisher, #BP510-500), 0.1 mg/mL BSA (BioShop, #ALB001.1), and 0.1% Triton-X 100

(Calbiochem, #9410-OP), pH7.4. RBD-HuFc was captured on AHC biosensors for 10 min at 5 µg/mL followed by 60 s in running buffer. RBD-HuFc captured AHC biosensors were placed in wells containing minibinders at concentrations ranging from 0.31 to 20 nM for 10 min, followed by 20 min of dissociation in running buffer. Prior to sol-RBD affinity measurements, Amine Reactive 2nd Generation (AR2G) biosensors (Sartorius, #18-5094) were pre-hydrated for 10 min in water before being activated using EDC/NHS chemistry (solution containing 20 mM EDC and 10 mM NHS in ddH2O) for 5 min. Sol-RBD in 10 mM Acetate buffer pH 5.0 was reacted with activated AR2G biosensors for 10 min and quenched for 5 min in 1M ethanolamine pH 8.5. Sol-RBD linked AR2G biosensors were equilibrated for 60 s in running buffer before measuring association of minibinders at concentrations ranging from 0.31 to 20 nM for 10 min, followed by 20 min of dissociation in running buffer. Minibinders were prepared in running buffer. For data evaluation, the ForteBio Data Analysis v11.0 software was used. The kinetic rate constants, association rate constant (ka, M-1s-1), dissociation rate constant (kd, s-1), and the equilibrium rate constant (KD, M) were determined by using a 1:1 Langmuir model.

<u>Multivalent Minibinder/S6P variant interaction kinetics via surface plasmon resonance (SPR)</u>
The Carterra LSA instrument was used to perform HT-SPRTM of minibinder affinity to Spike protein variants. To prepare the surfaces, the Single Flow Channel (SFC) and 96-Print Head (96PH) were primed with running buffer (HBS-T; 50 mM HEPES pH 7.5, 150 mM NaCl, 0.1% Tween 20). The capture surface was prepared in the 96PH by standard amine-coupling. A HC30-M chip (Carterra LSA cat# 4279) was activated with a 10-min injection of freshly prepared 1:1:1 (v/v/v) mixture of 0.4 M EDC + 0.1 M NHS + 0.1 M MES pH 5.5. Spike protein trimers were diluted to 12.5 µg/ml in 10 mM sodium acetate pH 4.5 (Carterra, #3628) and coupled for 20 min. Excess reactive esters were blocked with a 7-min injection of 1 M ethanolamine HCl pH 8.5 (Carterra, #3626). Final coupling levels were greater than 1000RU.

Minibinders were prepared in HBS-T buffer, at a three-fold dilution series for 6 points starting at 20nM. Association was for 20 minutes, with a 60-minute dissociation time. Samples were injected in ascending concentration without any regeneration. The data was double referenced in that both a local reference and a zero nanomolar analyte concentration (e.g. buffer) were subtracted. The double-referenced data were fit globally to a 1:1 Langmuir binding model in Carterra's Kinetic tool, allowing each spot its own ka and kd value to determine KD.

<u>Competition ELISA of minibinders and SARS-CoV-2 S-6P variants for immobilized hACE2-Fc</u>
0.003 mg/mL hACE2-Fc in 20mM Tris pH 8 and 100mM NaCl was immobilized to a Maxisorp 384-well plate (Thermo Scientific 464718) overnight at 4°C. Plates were slapped dry and blocked with Blocker Casein in TBS (Thermo Scientific 37532) for one hour at 37°C. 20nM mini binders were serially diluted 1:3 in avi-tagged prefusion-stabilized SARS-CoV-2 S6P (*23*) variants at their EC50 concentrations of 1.2nM for WT, 6.3nM for E406W, 1.7nM for K417N, 0.5nM for Y453F, 0.6nM for Y453R, 1.3nM for L455F, 1.1nM for F456L, 0.1nM for E484K, 0.2nM for N501Y, 0.3nM for B.1.1.7, 0.2nM for B.1.351, or 0.2nM for P.1 and incubated for 30 min on a non-binding plate (Greiner 781901) at 37°C. The plates with blocking buffer were slapped dry and the pre-incubated mini binders and spikes were added. Plates were incubated for 1 h at 37°C then washed 4x with TBST using a 405 TS Microplate Washer (BioTek) followed by addition of 30 µL avi-tag pAb (GenScript A00674) at 0.2 µg/mL. Plates were incubated for 1 h at 37°C then washed 4x with TBST using a 405 TS Microplate Washer (BioTek) followed by addition of 30 µL 1:2000 Goat

anti-Rabbit HRP (Invitrogen 656120) and a 1 h incubation at 37°C. Plates were washed 4x and TMB Microwell Peroxidase (Seracare 5120-0083) was added. The reaction was quenched after 2-3 min with 1 N HCl and the A450 of each well was read using a BioTek plate reader (BioTek). Data were plotted and fit in Prism (GraphPad) using nonlinear regression sigmoidal, 4PL, X is log(concentration) to determine IC50 values from curve fits. IC50 values less than two-fold below the concentration of S6P in that condition were not considered significantly different from two-fold below the concentration of S6P.

<u>Pseudovirus production</u>
HIV-based pseudotypes were prepared as previously described (*80*). Briefly, HEK293T cells were co-transfected using Lipofectamine 2000 (Life Technologies) with an S-encoding plasmid with full suite of variant mutations, an HIV Gag-Pol, Tat, Rev1B packaging construct, and the HIV transfer vector encoding a luciferase reporter according to the manufacturer's instructions. Variants produced include Wuhan-1, B.1.1.7, B.1.351, P.1, E406W, L452R, and Y453F. Cells were washed 3x with Opti-MEM and incubated for 5 h at 37°C with transfection medium. DMEM containing 10% FBS was added for 60 h. The supernatants were harvested by spinning at 2,500 x *g*, filtered through a 0.45 µm filter, concentrated with a 100 kDa membrane for 10 min at 2,500 x *g* and then aliquoted and stored at −80°C.

<u>Pseudovirus neutralization</u>
HEK-hACE2 cells were cultured in DMEM with 10% FBS (Hyclone) and 1% PenStrep with 8% $CO_2$ in a 37°C incubator (ThermoFisher). One day prior to infection, 40 µL of poly-lysine (Sigma) was placed into 96-well plates and incubated with rotation for 5 min. Poly-lysine was removed, plates were dried for 5 min then washed 1x with water prior to plating with 40,000 cells. The following day, cells were checked to be at 80% confluence. In an 80 µL final volume, minibinders were serially diluted in DMEM 1:3 starting at 100 nM. Pseudovirus was added 1:1 to the diluted minibinders and allowed to incubate for 30-60 min at room temperature. After incubation, the minibinder-virus mixture was added to the cells at 37°C and allowed to incubate for 2 h. Post infection, 160 µL of 20% FBS-2% PenStrep DMEM was added. After 48 h, 40 µL/well of One-Glo-EX substrate (Promega) was added to the cells and incubated in the dark for 5-10 min prior reading on a BioTek plate reader. Measurements were done in at least duplicate. Relative luciferase units were plotted and normalized in Prism (GraphPad). Nonlinear regression of log(inhibitor) versus normalized response was used to determine IC50 values from curve fits.

<u>SARS-CoV-2 neutralization</u>
Serial dilutions of minibinders were incubated with $10^2$ focus-forming units (FFU) of SARS-CoV-2 for 1 h at 37°C. Binder-virus complexes were added to Vero-hACE2-TMPRSS2 cell monolayers in 96-well plates and incubated at 37°C for 1 h. Subsequently, cells were overlaid with 1% (w/v) methylcellulose in MEM supplemented with 2% FBS. Plates were harvested 20-24 h later by removing overlays and fixed with 4% PFA in PBS for 20 min at room temperature. Plates were washed and sequentially incubated with an oligoclonal pool of SARS2-2, SARS2-11, SARS2-16, SARS2-31, SARS2-38, SARS2-57, and SARS2-71 anti-spike protein antibodies (*81*) and HRP-conjugated goat anti-mouse IgG in PBS supplemented with 0.1% saponin and 0.1% bovine serum albumin. SARS-CoV-2-infected cell foci were visualized using TrueBlue peroxidase substrate (KPL) and quantitated on an ImmunoSpot microanalyzer (Cellular Technologies). Data were processed using Prism software (GraphPad Prism 8.0).

Kidney organoid differentiation and infection with SARS-CoV-2

Kidney organoids were differentiated from H9 human embryonic stem cells (WiCell) for 21 days prior to infection in adherent, thin-layer Matrigel sandwich cultures induced for 36 hours with CHIR99021 as described previously (*41*). SARS-CoV-2 mutant strain B.1.351-HV001 containing E484K/N501Y/D614G mutations along with furin cleavage site point mutation were obtained directly from B.1.351 clinical isolates. All experiments using live viruses were performed at Biosafety Level 3 (BSL-3) facilities at the University of Washington in compliance with BSL-3 laboratory safety protocols (CDC BMBL 5th ed.) and the recent CDC guidelines for handling SARS-CoV-2. Virus stocks were generated and titrated using plaque forming assays in VERO cells (USAMRIID). Mini binders: FUS231-10GS, TRI2-2, and MON1 at 0.3 μM were diluted in serum-free DMEM and pre-incubated with virus (10 multiplicity of infection) for 1 h at 37°C. The virus: mini binder mix was then added to kidney organoids for 1 h at 37°C. After 1 h, kidney organoids were washed with 1X PBS and fresh organoid growth media (Advanced RPMI + 1X Glutamax + 1X B27 Supplement, Thermo Fisher Scientific) was added and incubated for 72 h. Supernatants were harvested from the infected organoids for plaque forming assays and cells were lysed using Trizol RNA extract for gene expression analysis.

Kidney organoid plaque assay

VERO cells were plated at 80% confluency, washed with 1X PBS, and incubated with serially diluted supernatant from infected organoids for 1 h at 37°C. Cells were then overlaid with a 1:1 mixture of 1.8% cellulose in water:2X DMEM supplemented with 4% heat-inactivated FBS, L-glutamine, 1X antibiotic-antimycotic (Gibco), and 220 mg/mL sodium pyruvate was layered on top of the cells and incubated for 48 h at 37°C. Cells were then fixed using 10% formaldehyde and incubated for 30 min at room temperature. Overlay was removed carefully and cells were stained with 0.5% crystal violet solution in 20% ethanol. Plaques were counted and the virus titer in the original sample was calculated as plaque-forming units per mL (PFU/mL). Data was analyzed using GraphPad Prism (8.0).

Kidney organoid gene expression analysis

RNA from infected kidney organoids was harvested using Trizol reagent. cDNA was generated using iScript™ cDNA Synthesis Kit (catalog #1708890, Bio-rad). qRT-PCR was performed using SYBR green master mix (catalog #4309155, Applied biosystems) and the following primers: SARS-CoV2-E F (GAACCGACGACGACTACTAGC), SARS-CoV2-E R (ATTGCAGCAGTACGCACACA), β-ACTIN F (GCAAAGACCTGTACGCCAACA), β-ACTIN R (ACACGGAGTACTTGCGCTCAG).

Selection of escape mutants in SARS-CoV-2 S using VSV-SARS-CoV-2 chimera

We used VSV-SARS-CoV-2 chimera (S from D614G strain) to select for SARS-CoV-2 S variants that escape peptide binder neutralization described as previously (*2*, *55*). Minibinder neutralization resistant mutants were recovered by plaque isolation. Briefly, plaque assays were performed to isolate the VSV-SARS-CoV-2 escape mutant on Vero cells with the indicated peptide binder in the overlay. The concentration of minibinder in the overlay was determined by neutralization assays at a multiplicity of infection (MOI) of 100. Escape clones were plaque-purified on Vero cells in the presence of peptide binder, and plaques in agarose plugs were amplified on MA104 cells with the peptide binder present in the medium. Viral stocks were amplified on MA104 cells at an MOI of 0.01 in Medium 199 containing 2% FBS and 20 mM HEPES pH 7.7 (Millipore Sigma) at 34°C. Viral supernatants were harvested upon extensive cytopathic effect and clarified of cell debris by centrifugation at 1,000 x *g* for 5 min. Aliquots were maintained at -80°C. Viral RNA was extracted from VSV-SARS-CoV-2 mutant viruses using RNeasy Mini kit (Qiagen), and S was amplified using OneStep RT-PCR Kit (Qiagen). The mutations were identified by Sanger sequencing (GENEWIZ).

Protein production for animal studies

Protein was produced by fermentation in the E. coli BL21 pLysS strain using pET vectors induced with IPTG. The 6xHis tagged proteins were purified from clarified cell lysates by immobilized metal chelate chromatography (IMAC, Ni-NTA resin) and step eluted with 300 mM imidazole. Proteins were polished by size exclusion chromatography using an S75 Increase Column into a final buffer of 20mM NaPO4 150mM NaCl pH 7.4. Proteins analyzed by SDS-PAGE after heating 95°C without (-) reducing agent Dithiothreitol (DTT) followed by Coomassie blue staining (molecular weight standards shown). Protein endotoxin levels are < 10 E.U./mg.

Mouse studies

Animal studies were carried out in accordance with the recommendations in the Guide for the Care and Use of Laboratory Animals of the National Institutes of Health. The protocols were approved by the Institutional Animal Care and Use Committee at the Washington University School of Medicine (assurance number A3381–01). Virus inoculations were performed under anesthesia that was induced and maintained with ketamine hydrochloride and xylazine, and all efforts were made to minimize animal suffering.

Heterozygous K18-hACE C57BL/6J female mice (strain: 2B6.Cg-Tg(K18-ACE2)2Prlmn/J) were obtained from The Jackson Laboratory. Animals were housed in groups and fed standard chow diets. For inoculation, 8-week-old mice were administered $10^3$ PFU of the indicated SARS-CoV-2 strain via intranasal administration. The previously described influenza minibinder was used as a negative control (*82*).

Measurement of viral burden in mouse studies

Tissues were weighed and homogenized with zirconia beads in a MagNA Lyser instrument (Roche Life Science) in 1 mL of DMEM media supplemented with 2% heat-inactivated FBS. Tissue homogenates were clarified by centrifugation at 10,000 rpm for 5 min and stored at −80°C. RNA was extracted using the MagMax mirVana Total RNA isolation kit (Thermo Scientific) on a Kingfisher Flex extraction robot (Thermo Scientific). RNA was reverse transcribed and amplified using the TaqMan RNA-to-CT 1-Step Kit (ThermoFisher). Reverse transcription was performed at 48°C for 15 min followed by 2 min at 95°C. Amplification was accomplished over 50 cycles as follows: 95°C for 15 s and 60°C for 1 min. Copies of SARS-CoV-2 N gene RNA in samples were determined using a previously published assay (*42*, *83*). Briefly, a TaqMan assay was designed to target a highly conserved region of the N gene (Forward primer: ATGCTGCAATCGTGCTACAA; Reverse primer: GACTGCCGCCTCTGCTC; Probe: /56-FAM/TCAAGGAAC/ZEN/AACATTGCCAA/3IABkFQ/). This region was included in an RNA standard to allow for copy number determination down to 10 copies per reaction. The reaction mixture contained final concentrations of primers and probe of 500 and 100 nM, respectively.

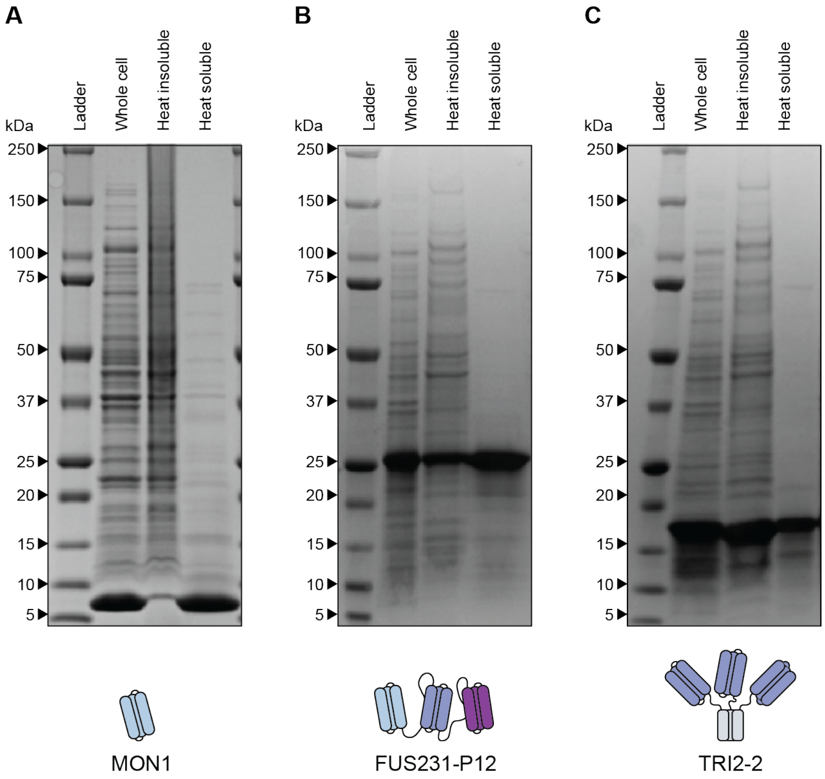

**Fig. S1.**
Heat-based purification of mono and multivalent minibinders. Proteins were expressed in E. coli using standard protein overexpression procedures. Harvested cells were heat treated at 70 °C for 5 min to lyse cells and precipitate cellular proteins. (**A**) SDS PAGE of MON1. (**B**) SDS PAGE of FUS231-P12. (**C**) SDS PAGE of TRI2-2.

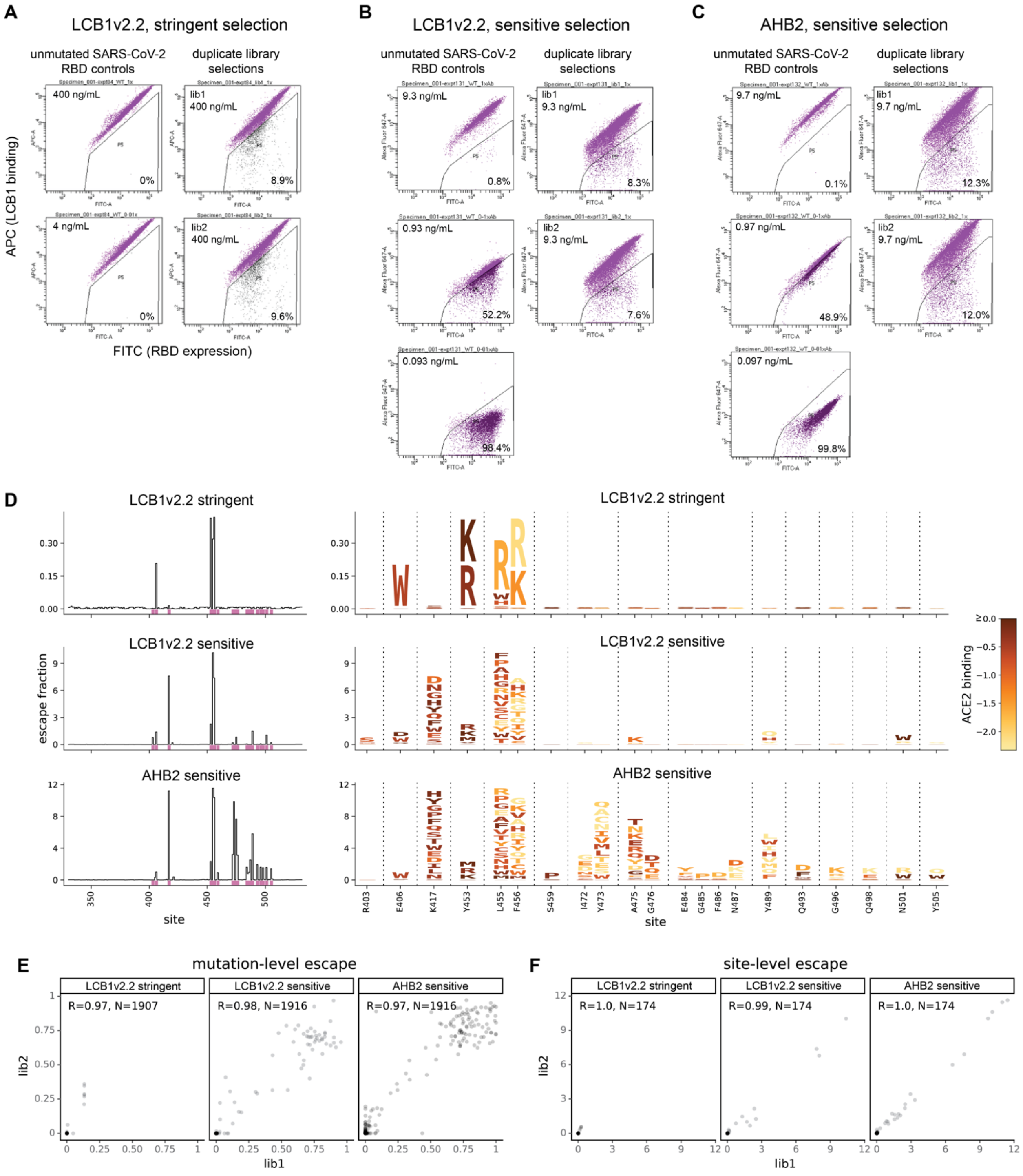

**Fig. S2.**
Site saturation mutagenesis of RBD to predict MON1 (LCB1v2.2) and MON2 (AHB2) escape mutants. (**A, B, C**) FACS gates for deep mutational scanning selections. Duplicate yeast display libraries expressing all mutations tolerated in the SARS-CoV-2 RBD were labeled with LCB1v2.2 (**A, B**) or AHB2 (**C**). The population of cells indicated escaping minibinder binding was sorted and deep sequenced, enabling calculation of the "escape fraction" of each RBD mutation (the fraction of yeast expressing that mutant that were found in the escape bin). (**D**) Miniprotein escape

profiles averaged across duplicate deep mutational scanning library selections. Line plots, left, show the total escape (sum of per-mutation escape) at all sites in the RBD. Sites marked in pink are shown in logoplots, right. Logoplots illustrate per-mutation escape by the height of letters reflecting individual amino acid mutations. Letters are colored according to their previously measured effects on ACE2-binding affinity (*37*), according to the scale bar at th right. (**E, F**) Correlation in escape between duplicate libraries, at the level of individual mutations (**E**) or total escape per site (**F**).

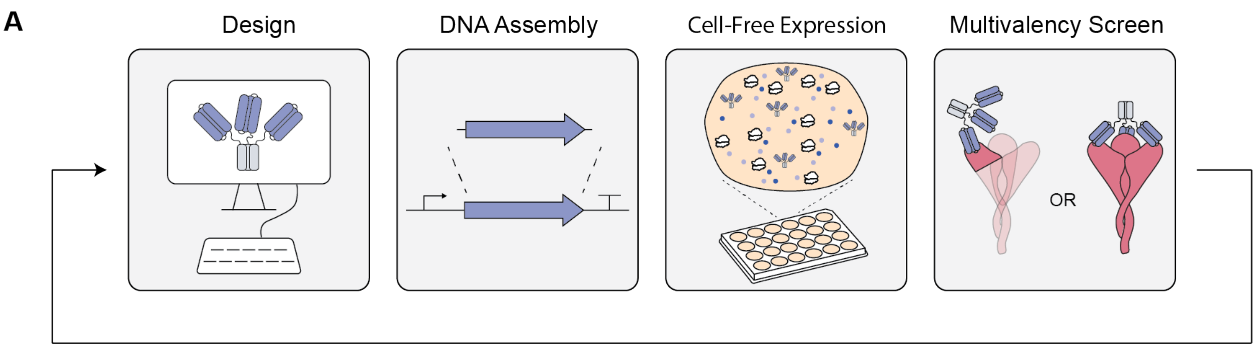

**B** **Cell-based protein expression**

| Day | Step |
|---|---|
| 1 | Gibson assembly |
| 1 | Transformation |
| 2 | Pick colonies for miniprep |
| 3 | Miniprep and sequencing |
| 4 | Sequencing results |
| 4 | Transform expression strain |
| 5 | Start expression culture |
| 6 | Harvest expression culture |
| 6 | Cell lysis |
| 6 | Protein purification |
| 6 | Extensive protein dialysis |
| 7 | Use protein |

**C** **Cell-free protein expression**

| Day | Step |
|---|---|
| 1 | Gibson assembly |
| 1 | LET PCR |
| 1 | CFPS |
| 2 | Protein purification |
| 2 | Extensive protein dialysis |
| 3 | Use protein |

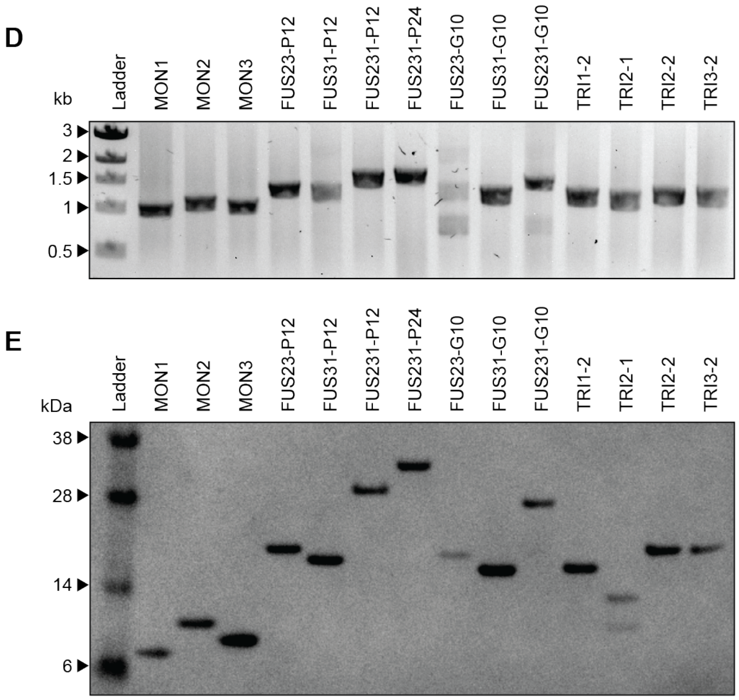

**Fig. S3**
Cell-free DNA assembly and protein synthesis of multivalent minibinders. (**A**) A cell-free workflow for the expression and evaluation of multivalent minibinders. (**B**) Step-by-step workflow for cell-based DNA assembly and protein expression. (**C**) Standard step-by-step workflow for cell-free DNA assembly and protein expression. (**D**) Agarose gel of linear DNA templates for CFPS assembled via Gibson assembly and amplified via PCR. (**E**) SDS PAGE of purified proteins expressed via CFPS.

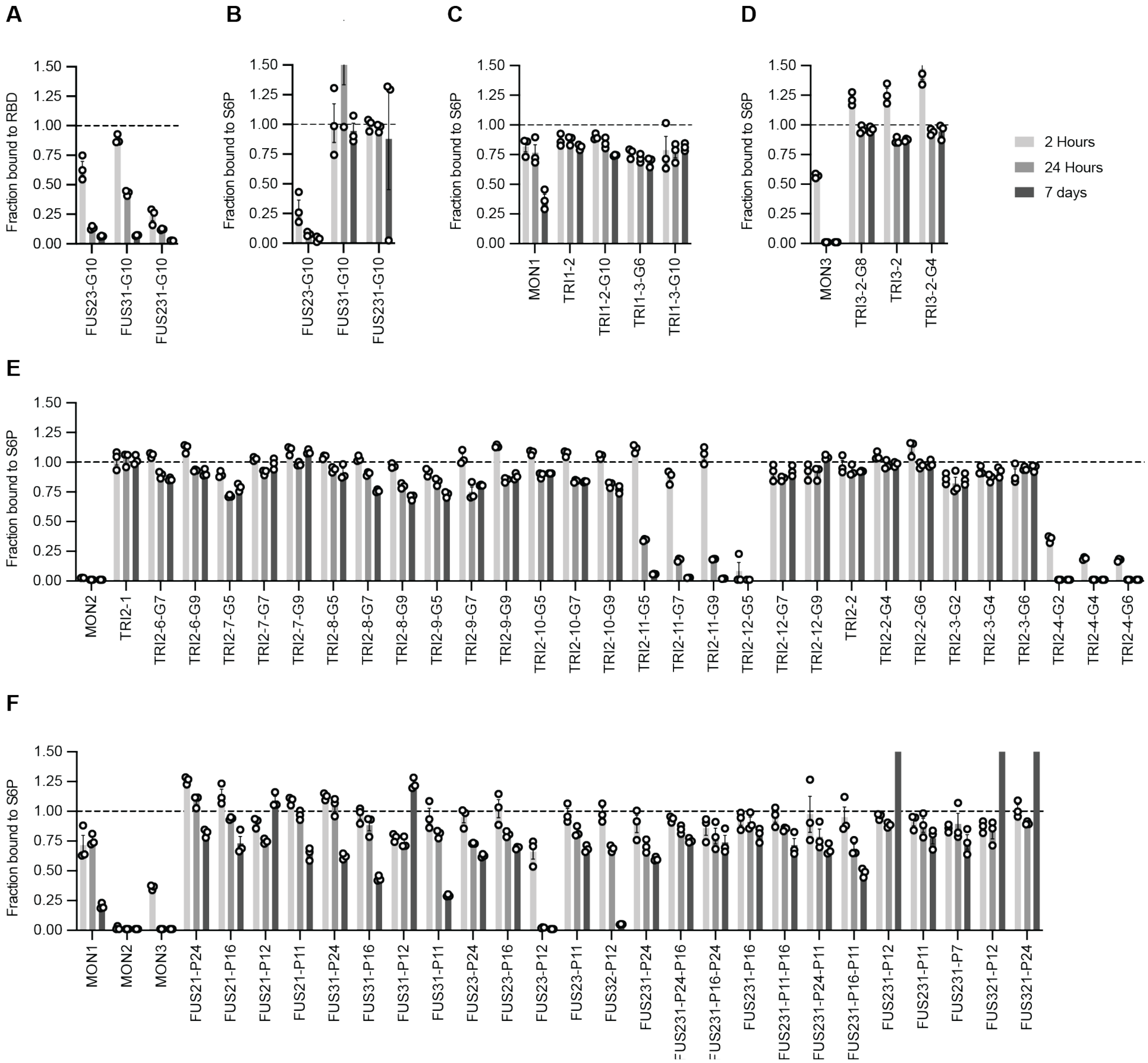

**Fig. S4.**
Off-rate screen of multivalent minibinders. Dissociation of minibinder constructs from RBD or S6P over time was measured via competition AlphaLISA. (mean ± SEM, n=3 replicates from a single experiment) (**A**) Dissociation of FUS proteins with G10 linkers from RBD measured via competition with MON1. (**B**) Dissociation of FUS proteins with G10 linkers from S6P measured via competition with MON1. (**C**) Dissociation of TRI1 proteins from S6P measured via competition with TRI2-1. (**D**) Dissociation of TRI3 proteins from S6P measured via competition with TRI2-1. (**E**) Dissociation of TRI2 proteins from S6P measured via competition with TRI2-1. (**F**) Dissociation of FUS proteins from S6P measured via competition with FUS231-P24.

A

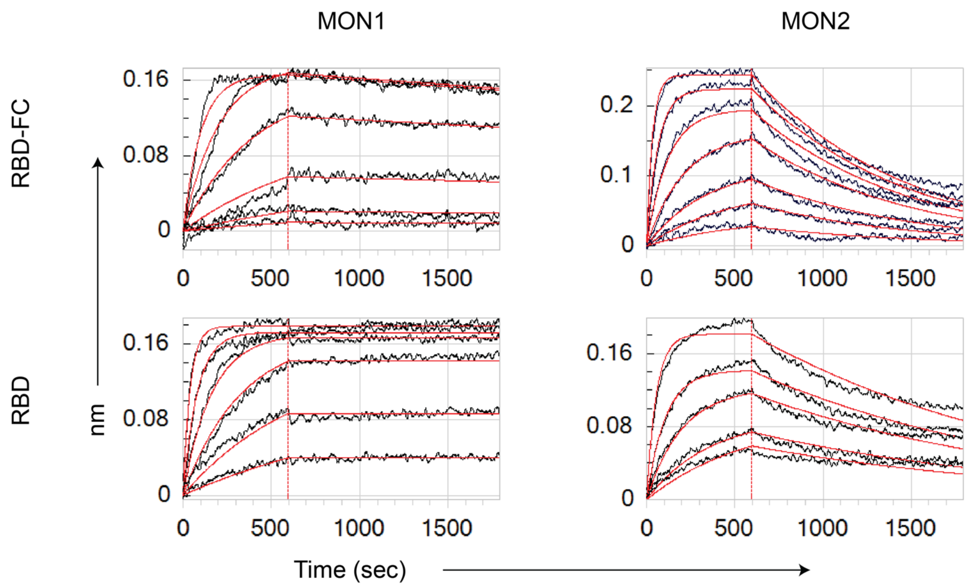

B

| | | MON1 | MON2 |
|---|---|---|---|
| RBD-FC | $k_a$ ($x10^5\ M^{-1}s^{-1}$) | 9.83 | 14.2 |
| | $k_d$ ($x10^{-5}\ s^{-1}$) | 8.41 | 114 |
| | $K_D$ ($x10^{-12}$ M) | 85.6 | 803 |
| RBD | $k_a$ ($x10^5\ M^{-1}s^{-1}$) | 12.5 | 17.4 |
| | $k_d$ ($x10^{-5}\ s^{-1}$) | <4.27 | 61.8 |
| | $K_D$ ($x10^{-12}$ M) | <34.2 | 355 |

**Fig. S5.**
Kinetics of MON1, and MON2 binding to the SARS-CoV-2 RBD. Minibinders were titrated from 20 to 0.31 nM against either Human FC tagged RBD (RBD-FC) captured by AHC biosensors or monovalent soluble RBD (RBD) covalently linked to AR2G biosensors. (**A**) Sensograms of minibinders binding to RBD. Red curves illustrated fits obtained from 1:1 Langmuir binding model. Sensorgram images were copied directly from export files from ForteBio Data Analysis v11.0 software without modification. (**B**) Table of determined kinetic and equilibrium binding parameters for each minibinder. The < symbol indicates a cut-off of less than 5% dissociation observed during the dissociation phase, indicating insufficient time to accurately quantify the dissociation rate constant.

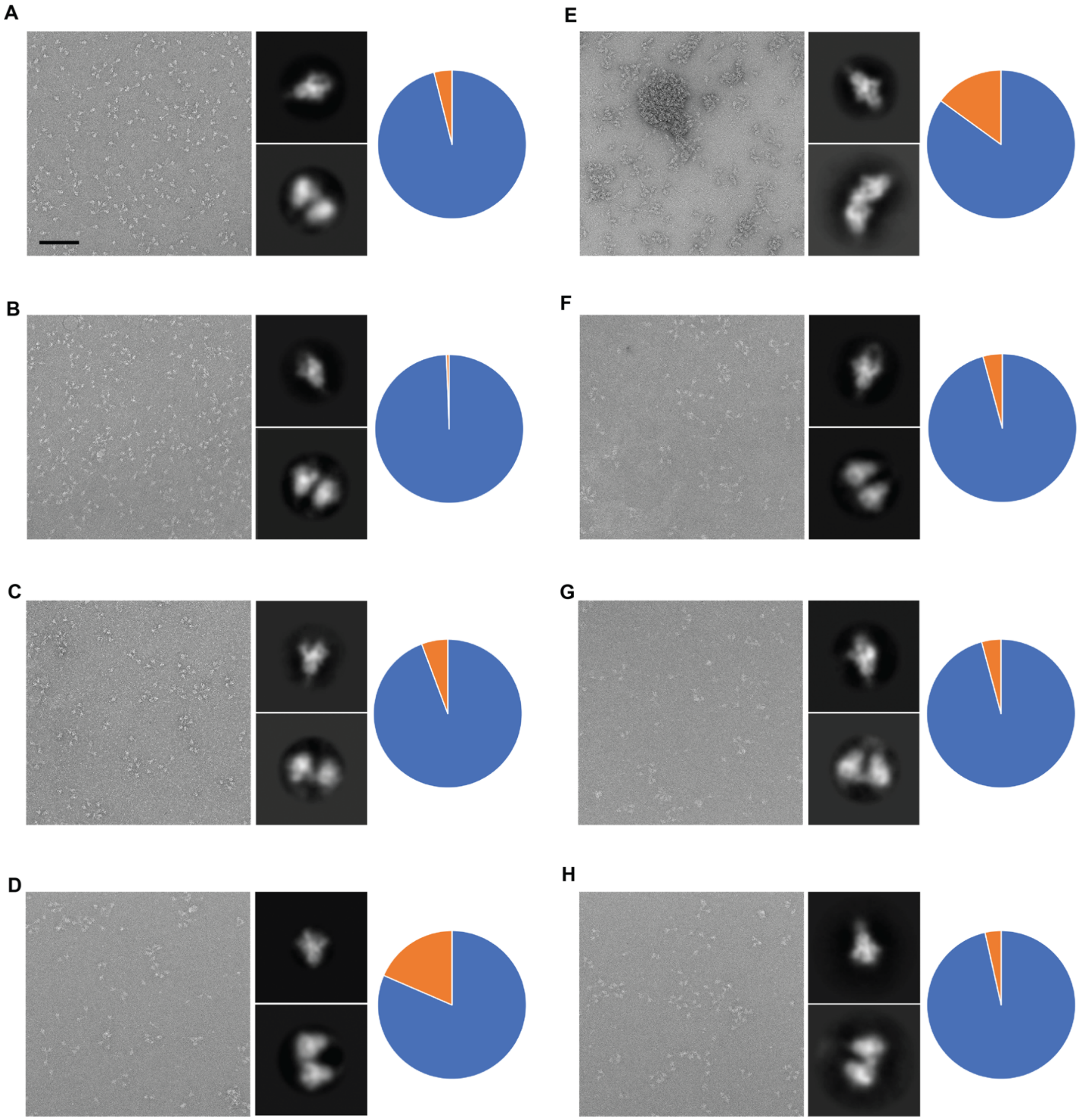

**Fig. S6.**
Negative stain EM analysis of minibinder-mediated crosslinking/aggregation of spike ectodomain trimers upon complex formation. Representative electron micrograph and 2D class averages of SARS-CoV-2 S in complex with TRI2-2 (**A**), FUS231-G10 (**B**), FUS231-P24 (**C**), FUS31-G8 (**D**), TRI1-5-G2 (**E**), TRI1-5-G4 (**F**), TRI1-5-G6 (**G**) or FUS31-G10 (**H**). Scale bar: 100 nm. After two-dimensional classification, the number of particles assigned to well-defined classes with single (blue) or two neighboring cross-linked (orange) spike trimers are presented as pie charts. The fraction of total cross-linked trimers is underestimated since higher-order cross-linked trimers did not yield well-defined 2D averages.

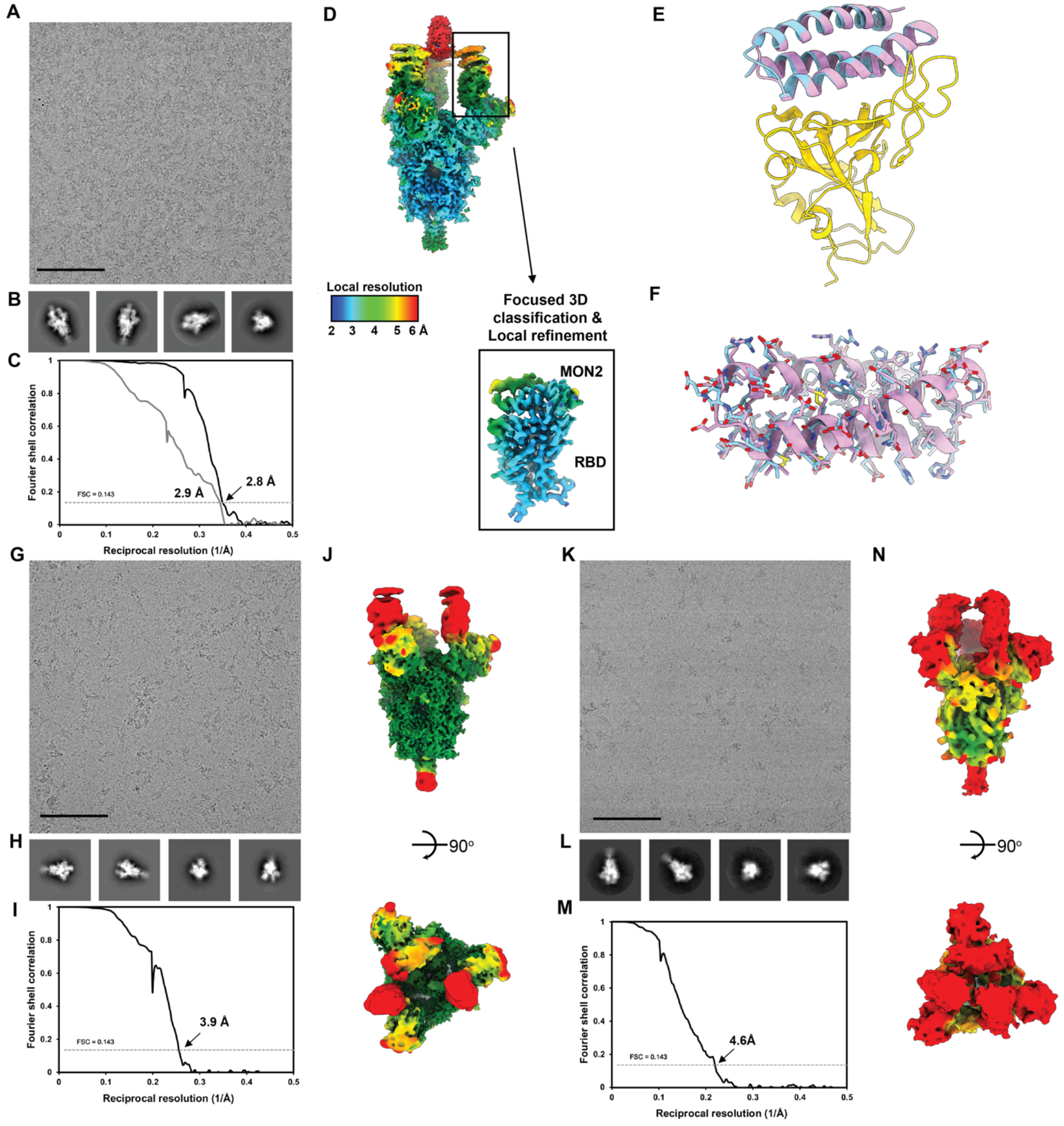

**Fig. S7.**

CryoEM data processing of the S6P/TRI2-2, S6P/FUS31-G10, and S6P/FUS231-P24 datasets. **(A-B)** Representative electron micrograph and 2D class averages of SARS-CoV-2 S in complex with TRI2-2 embedded in vitreous ice. Scale bar: 100 nm. (**C)** Gold-standard Fourier shell correlation curves for TRI2-2 bound SARS-CoV-2 S trimer (black line) and locally refined RBD/TRI2-2 domains (gray line). The 0.143 cutoff is indicated by a horizontal dashed line. (**D)** Local resolution maps calculated using cryoSPARC for the whole reconstruction as well as for the locally refined RBD/MON2 region. (**E)** Ribbon diagram of the RBD/MON2 designed model (pink) superimposed with the MON2 cryoEM structure (cyan) (**F**) MON2 designed model (pink) superimposed with the S6P/TRI2-2 cryoEM structure (pink) with side chains displayed as sticks. **(G-J)** Representative electron micrograph (**G**), 2D class averages (**H**), gold-standard Fourier shell correlation curve (**I**) and local resolution map (**J**) of SARS-CoV-2 S in complex with FUS231-P24 embedded in vitreous ice. **(K-N)** Representative electron micrograph (**K**), 2D class averages (**L**), gold-standard Fourier shell correlation curve (**M**), and local resolution map (**N**) of SARS-CoV-2 S in complex with FUS31-G10 embedded in vitreous ice.

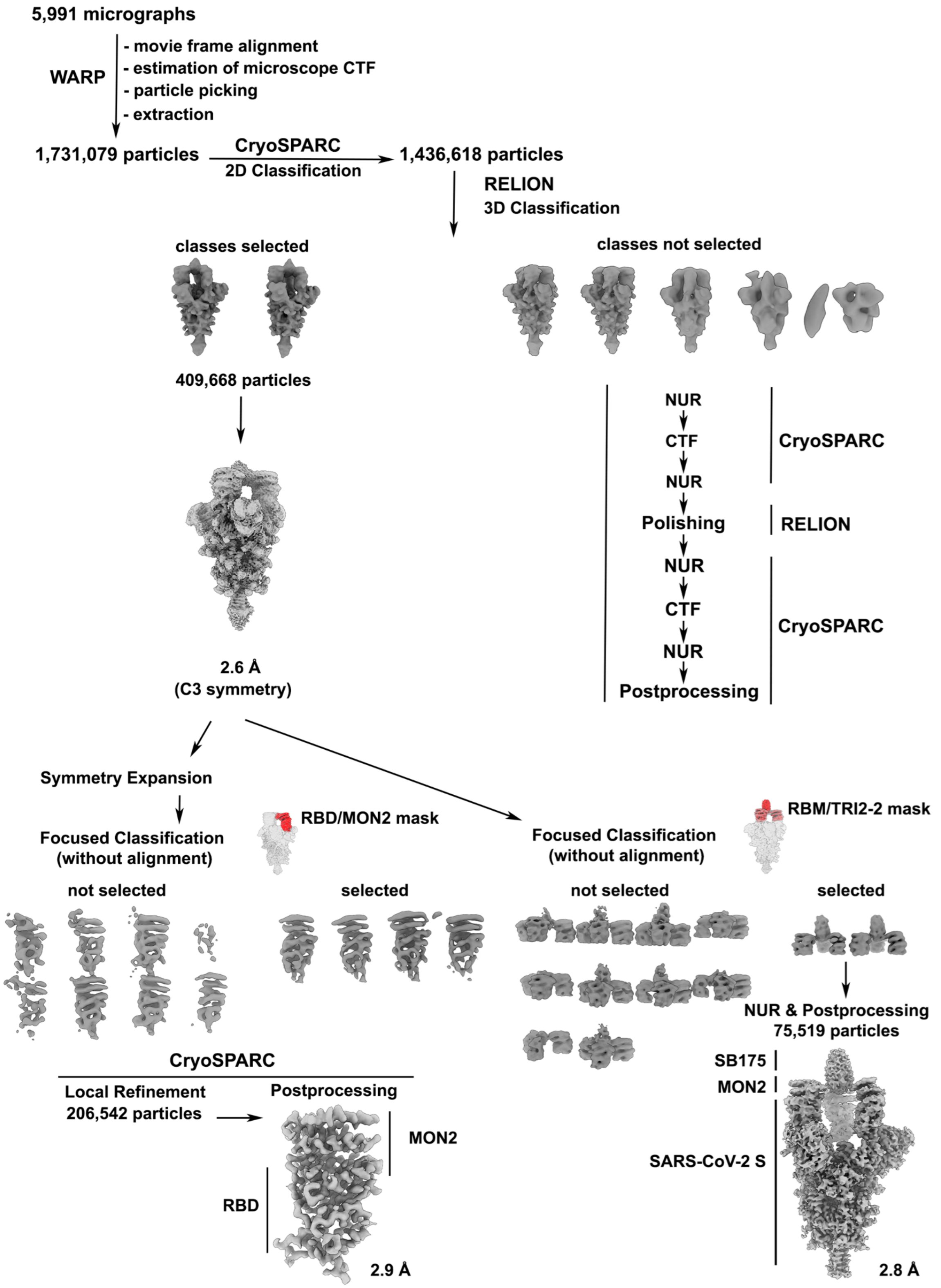

**Figure S8.**
CryoEM processing scheme of SARS-CoV-2 S/TRI2-2 complex.

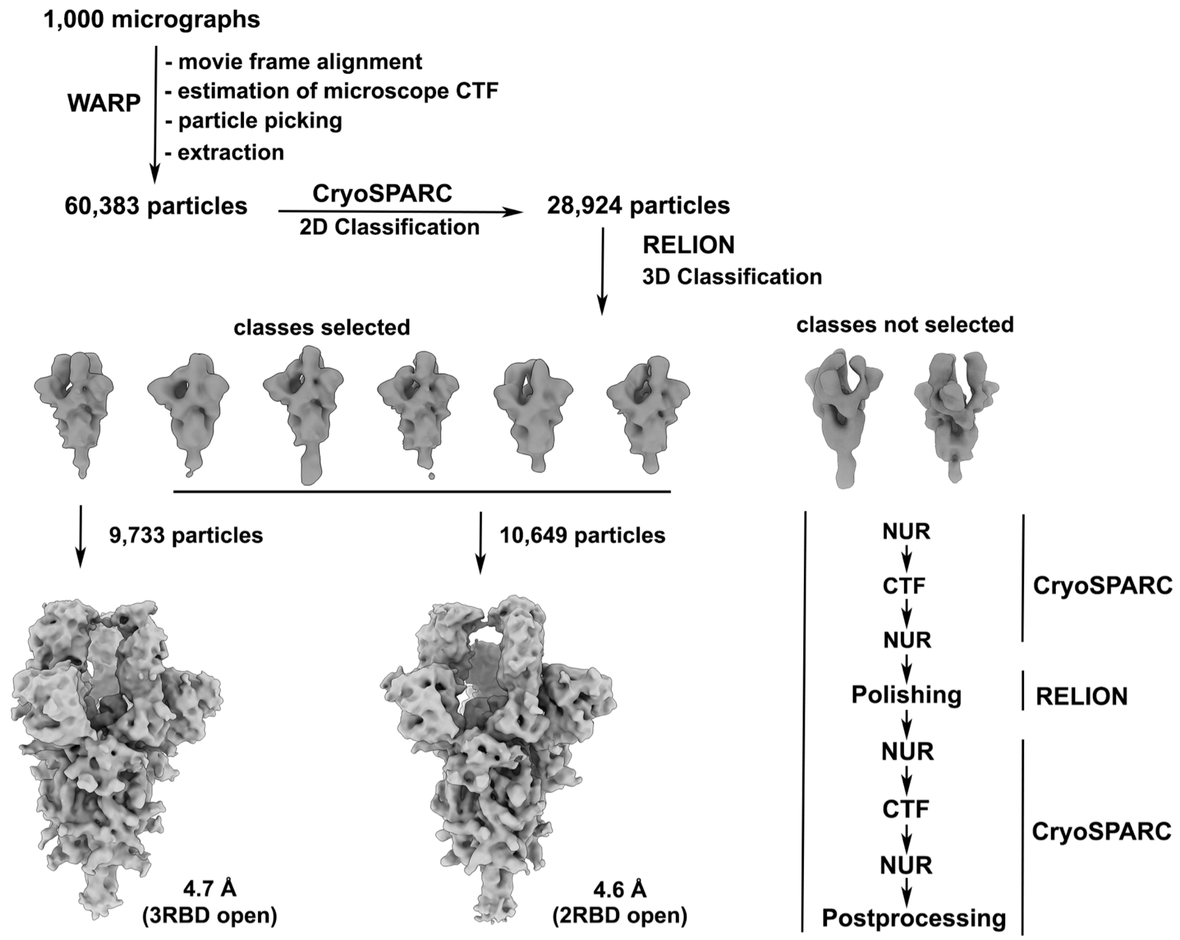

**Figure S9.**
CryoEM processing scheme of SARS-CoV-2 S/FUS31-G10 complex.

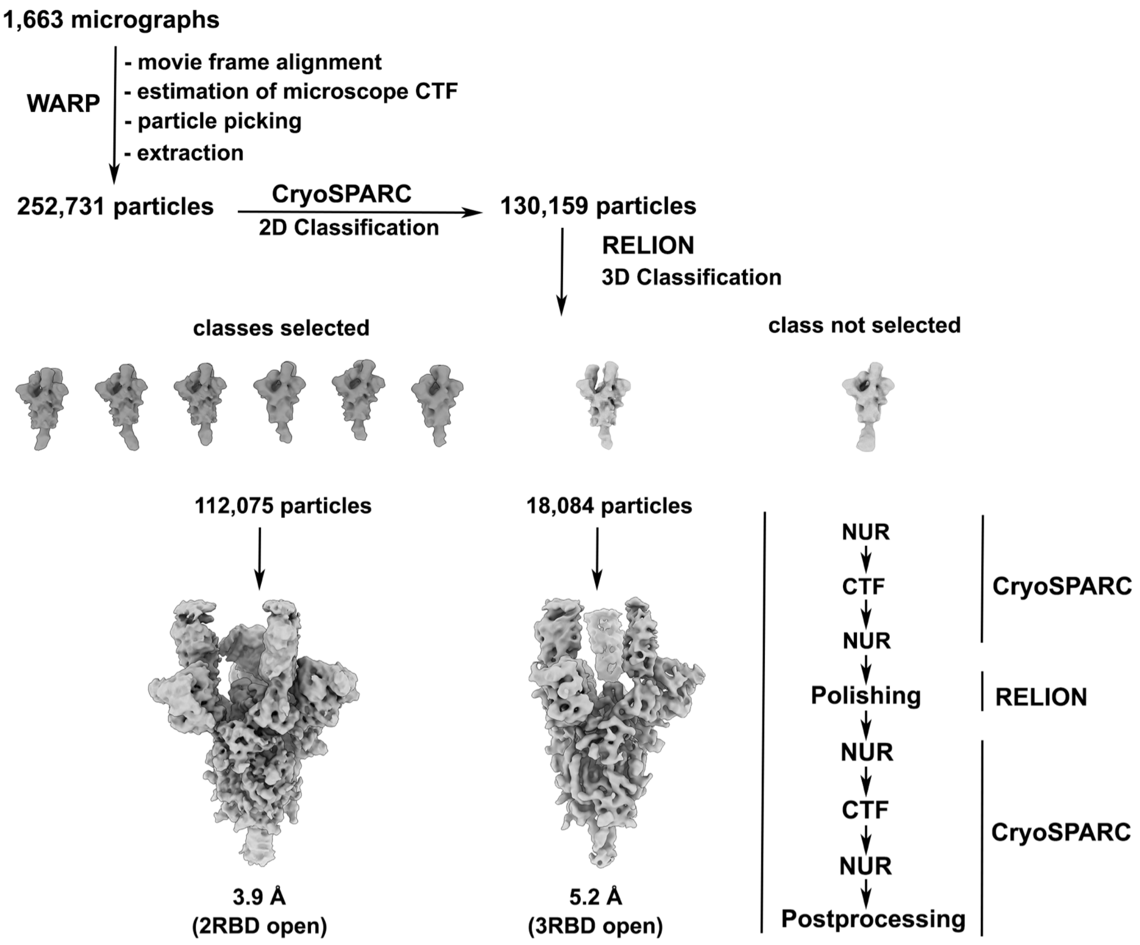

**Figure S10.**
CryoEM processing scheme of SARS-CoV-2 S/FUS231-P24 complex.

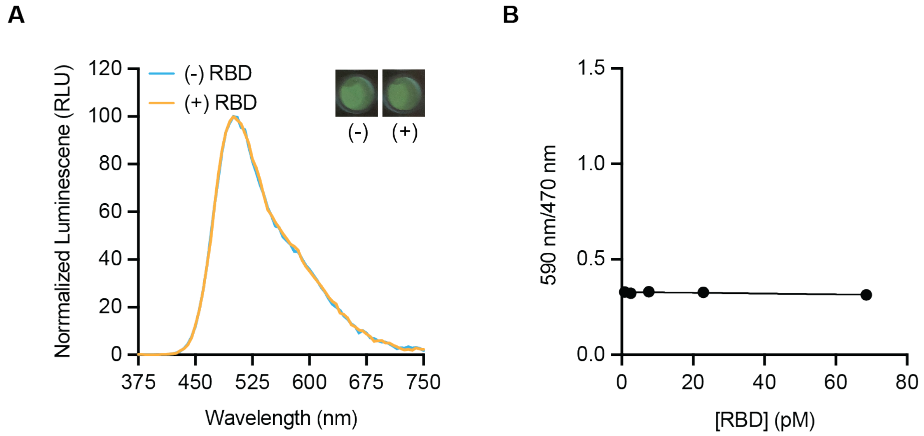

**Fig. S11.**
FUS231-P12 BRET sensor does not detect monomeric RBD. (**A**) Luminescence emission spectra and image of the BRET sensor (100 pM) in the presence (yellow trace, 100 pM) and absence (blue trace) of RBD. (**B**) Titration of RBD at 100 pM sensor protein (Mean ± SEM, n=3 replicates from a single experiment).

**A**

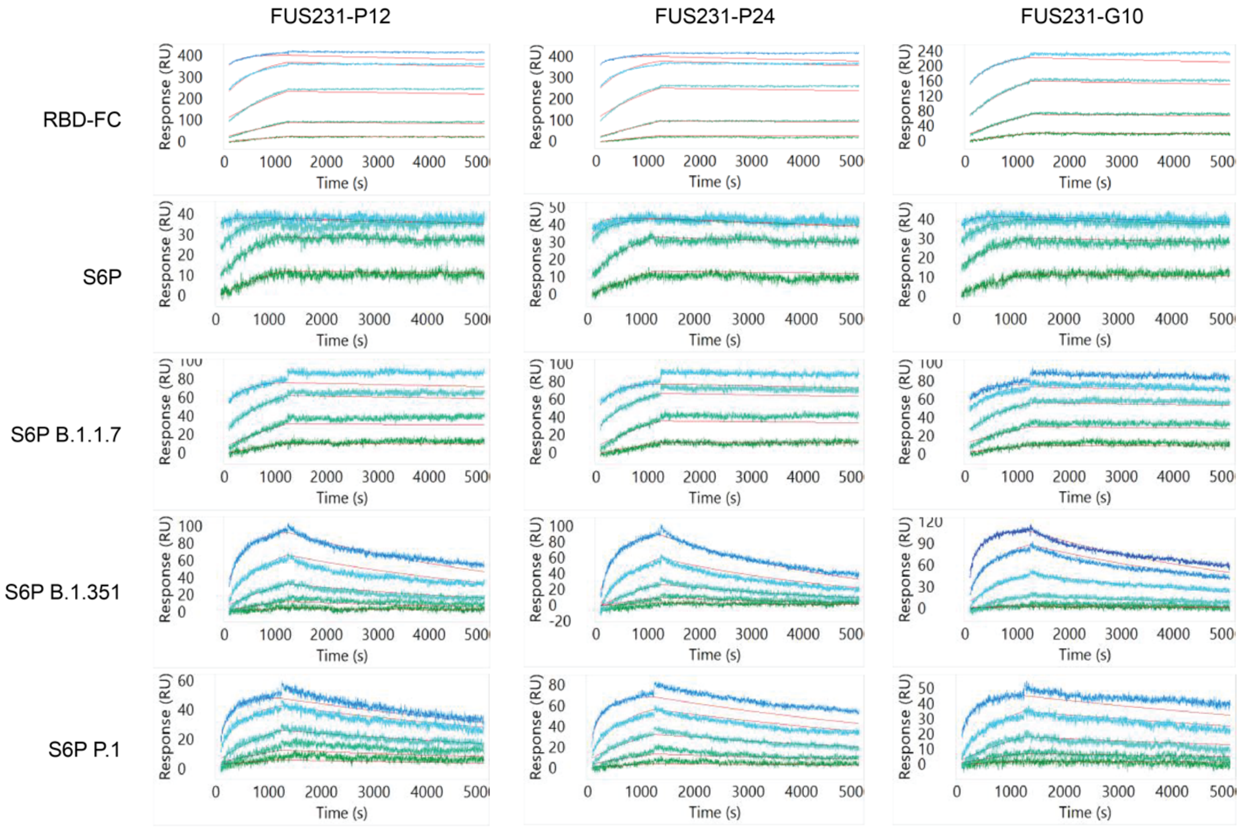

**B**

| | | FUS231-P12 | FUS231-P24 | FUS231-G10 |
|---|---|---|---|---|
| RBD-FC | $k_a$ (x$10^5$ $M^{-1}s^{-1}$) | 6.5 | 7.5 | 9.9 |
| | $k_d$ (x$10^{-5}$ $s^{-1}$) | <1.4 | <1.4 | <1.4 |
| | $K_D$ (x$10^{-12}$ M) | <21.9 | <18.8 | <14.4 |
| S6P | $k_a$ (x$10^5$ $M^{-1}s^{-1}$) | 38.2 | 38.9 | 29.2 |
| | $k_d$ (x$10^{-5}$ $s^{-1}$) | 1.7 | 2.7 | 1.9 |
| | $K_D$ (x$10^{-12}$ M) | 4.5 | 7.1 | 6.5 |
| S6P B.1.1.7 | $k_a$ (x$10^5$ $M^{-1}s^{-1}$) | 14.8 | 17.4 | 10.8 |
| | $k_d$ (x$10^{-5}$ $s^{-1}$) | <1.4 | <1.4 | <1.4 |
| | $K_D$ (x$10^{-12}$ M) | <9.6 | <82.1 | <13.2 |
| S6P B.1.351 | $k_a$ (x$10^5$ $M^{-1}s^{-1}$) | 4.2 | 4.3 | 2.2 |
| | $k_d$ (x$10^{-5}$ $s^{-1}$) | 17.6 | 25.8 | 19.9 |
| | $K_D$ (x$10^{-12}$ M) | 411.0 | 597.4 | 899.8 |
| S6P P.1 | $k_a$ (x$10^5$ $M^{-1}s^{-1}$) | 8.1 | 6.2 | 5.3 |
| | $k_d$ (x$10^{-5}$ $s^{-1}$) | 10.9 | 12.2 | 8.6 |
| | $K_D$ (x$10^{-12}$ M) | 133.5 | 195.3 | 163.1 |

**Fig. S12.**
Kinetic analysis of interactions between multivalent minibinders and SARS-CoV-2 S protein variants. Multivalent minibinders were injected at concentrations ranging from 20 nM to 0.08 nM in three-fold serial dilutions against S protein variants covalently coupled to the chip via amine coupling. The double-referenced data were fit globally to a simple 1:1 Langmuir binding model in Carterra's Kinetic tool. (**A**) Sensograms of FUS231 proteins binding to the RBD, S6P, and S6P mutants. (**B**) Table of kinetic and equilibrium parameters of the measured interactions.

A

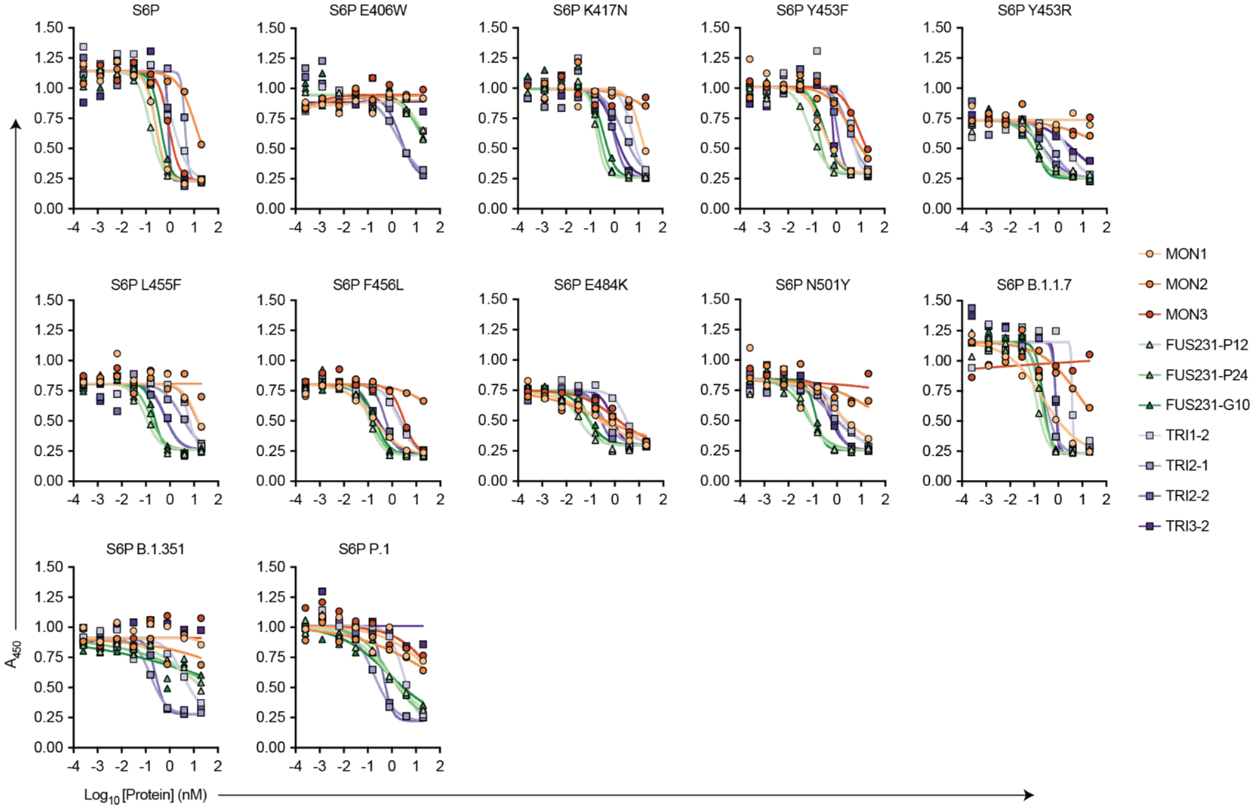

B

| Protein | ACE Competition ELISA $IC_{50}$ (pM) | | | | | | | | | | | |
|---|---|---|---|---|---|---|---|---|---|---|---|---|
| | WT | E406W | K417N | Y453F | Y453R | L455F | Y456L | E484K | N510Y | B.1.1.7 | B.1.351 | P.1 |
| MON1 | 261* | N/A | 12,250 | 286 | N/A | 12,430 | 181* | 167 | 1,403 | 225 | N/A | N/A |
| MON2 | 11,700 | N/A | N/A | 4,011 | N/A | N/A | N/A | 237 | N/A | 8,482 | N/A | N/A |
| MON3 | 896 | N/A | N/A | 8,743 | N/A | 11,870 | 3,079 | 1,028 | N/A | N/A | N/A | N/A |
| FUS231-P12 | 153* | N/A | 191* | 83* | 225* | 95* | 111* | 31* | 67* | 118* | 6,525 | 727 |
| FUS231-P24 | 260* | 18,300 | 245* | 175* | 96* | 156* | 134* | 81 | 55* | 187 | 12,820 | 803 |
| FUS231-G10 | 415* | 18,930 | 361* | 319 | 138* | 223* | 194* | 74 | 132 | 240 | N/A | 733 |
| TRI1-2 | 1,388 | N/A | 5,628 | 4,479 | 2,458 | 5,897 | 1,939 | 2,721 | 1,404 | 3,830 | 3,832 | 2,997 |
| TRI2-1 | 4,247 | 1,720* | 2,412 | 3,362 | 573 | 3,052 | 2,182 | 329 | 631 | 267 | 165 | 175 |
| TRI2-2 | 870 | 2,242* | 934 | 1,090 | 249* | 648* | 543* | 244 | 625 | 792 | 270 | 429 |
| TRI3-2 | 844 | N/A | 1086 | 722 | 3,936 | 627* | 246* | 772 | 464 | 786 | N/A | N/A |

C

| S6P Variant | [S6P] (nM) |
|---|---|
| WT | 1.2 |
| E406W | 6.9 |
| K417N | 1.7 |
| Y453F | 0.5 |
| Y453R | 0.6 |
| L455F | 1.3 |
| F456L | 1.1 |
| E484K | 0.1 |
| N501Y | 0.2 |
| B.1.1.7 | 0.3 |
| B.1.351 | 0.2 |
| P.1 | 0.2 |

D

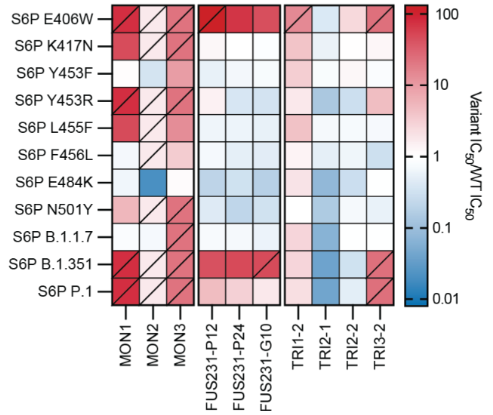

**Fig. S13.**

Competition of ACE2 and mini binder constructs for S6P. **(A)** Competition ELISA curves (mean, n=2). **(B)** Summary of minibinder construct competition $IC_{50}$ values. An * denotes an $IC_{50}$ value less than 2-fold lower than the limiting concentration of S6P variant present in the well. N/A indicates an $IC_{50}$ value higher than the tested concentration range and greater than 20,000 pM. **(C)** $EC_{50}$ values of S6P variants binding to ACE2 used as the concentration for minibinder construct competition. **(D)** Ratio of mutant to wild type (WT) $IC_{50}$ values of minibinder constructs. The ratio in cells containing a slash was determined using the highest tested minibinder construct concentration.

A

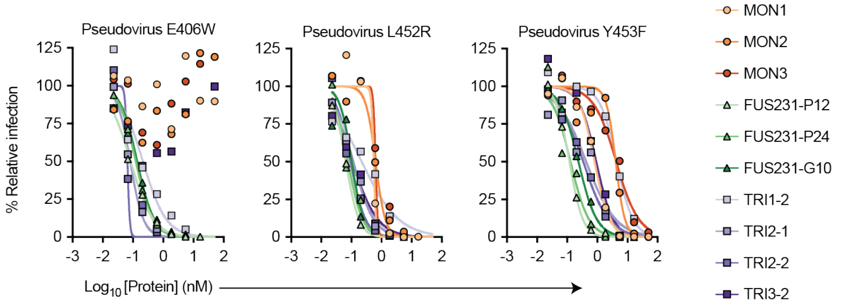

B

| Protein | HIV Pseudovirus Neutralization $IC_{50}$ (pM) | | |
|---|---|---|---|
| | E406W | L452R | Y453F |
| MON1 | N/A | 570 | 817 |
| MON2 | N/A | 639 | 4,102 |
| MON3 | N/A | 637 | 4,260 |
| FUS231-P12 | 84 | 65 | 120 |
| FUS231-P24 | 124 | 77 | 130 |
| FUS231-G10 | 138 | 103 | 237 |
| TRI1-2 | 214 | 223 | 4,406 |
| TRI2-1 | 81 | 117 | 430 |
| TRI2-2 | 67 | 94 | 343 |
| TRI3-2 | N/A | 109 | 928 |

**Fig. S14.**

Pseudovirus neutralization of additional SARS-CoV-2 spike point mutations. (**A**) Neutralization of SARS-CoV-2 pseudovirus variants by minibinder constructs (mean, n=2). (**B**) Table summarizing neutralization potencies of multivalent minibinder constructs against SARS-CoV-2 pseudovirus variants. N/A indicates an $IC_{50}$ value above the tested concentration range and an $IC_{50}$ greater than 50,000 pM.

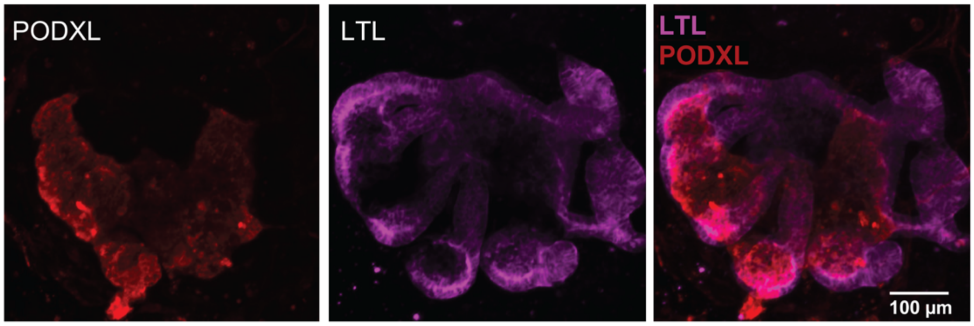

**Fig. S15.**
Representative confocal images of a human kidney organoid derived from H9 human embryonic stem cells (LTL, *Lotus tetragonolobus* lectin, proximal tubule marker in magenta; PODXL, podocalyxin, podocyte marker in red).

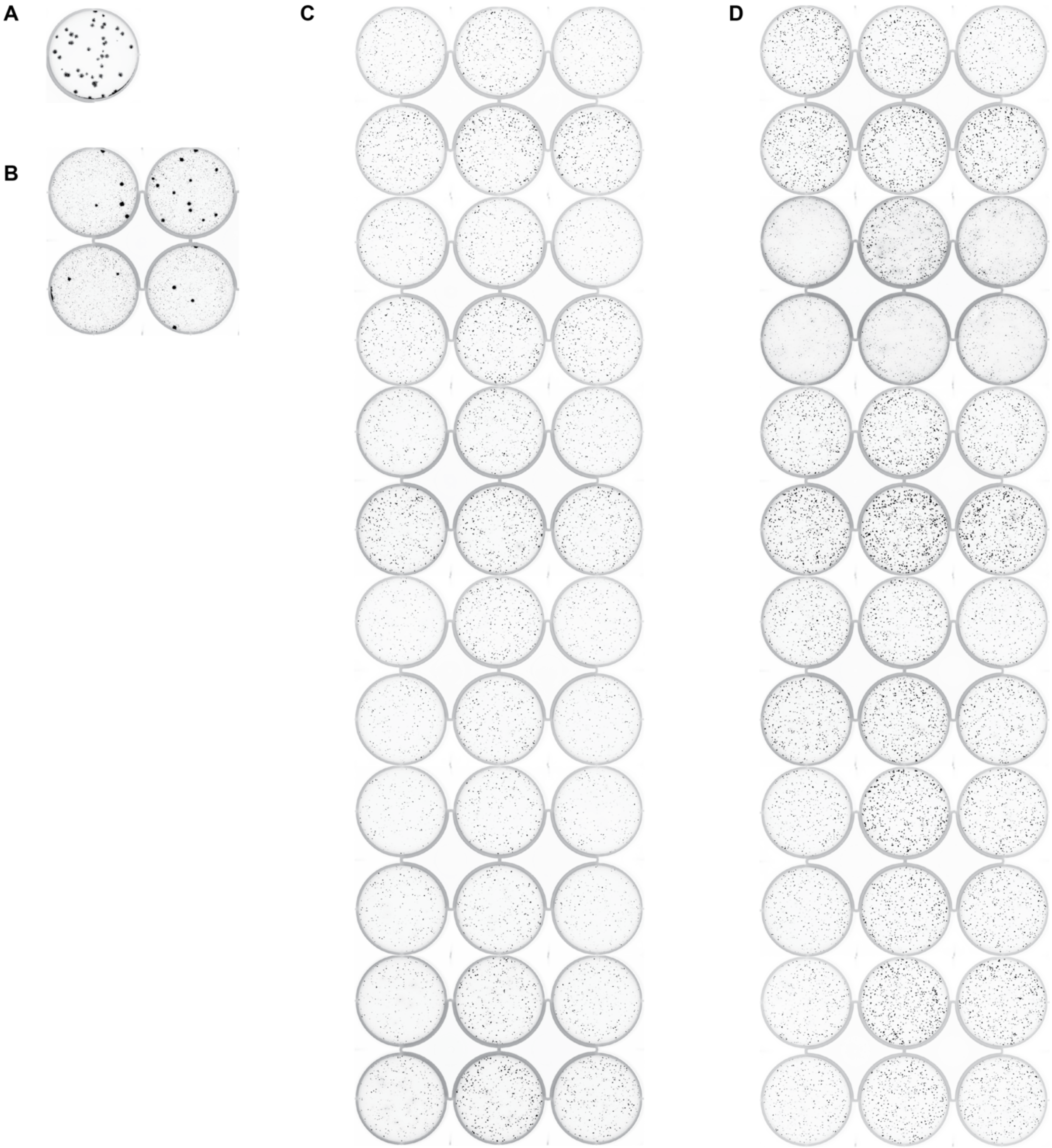

**Fig. S16.**
Plaque assays were performed to isolate VSV-SARS-CoV-2 chimera virus escape mutants against a control neutralizing antibody (2B04) and the FUS231-P12 and TRI2-2 multivalent minibinders. Large plaques are indicative of escape. FUS231-P12 and TRI2-2 replicates were performed in three separate experiments consisting of two plates each. (**A**) No inhibitor in overlay. (**B**) 2B04 neutralizing mAb in overlay. (**C**) FUS231-P12 in overlay. (**D**) TRI2-1 in overlay.

**Table S1.**
List of abbreviations used to describe multivalent minibinders in this paper.

| Abbreviation | Protein (N to C) |
|---|---|
| MON1 | LCB1v2.2 |
| MON2 | AHB2 |
| MON3 | LCB3v2.2 |
| FUS23-P12 | AHB2v2-PAS12-LCB3v2.2 |
| FUS31-P12 | LCB3v2.2-PAS12-LCB1v2.2 |
| FUS231-P12 | AHB2v2-PAS12-LCB3v2.2-PAS12-LCB1v2.2 |
| FUS231-P24 | AHB2v2-PAS24-LCB3v2.2-PAS24-LCB1v2.2 |
| FUS23-G10 | AHB2v2-GS10-LCB3v2.2 |
| FUS31-G10 | LCB3v2.2-GS10-LCB1v2.2 |
| FUS231-G10 | AHB2v2-GS10-LCB3v2.2-GS10-LCB1v2.2 |
| TRI1-2 | SB175-6GS-LCB1v2.2 |
| TRI2-1 | AHB2-4GS-1rfo |
| TRI2-2 | AHB2-2GS-SB175 |
| TRI3-2 | LCB3v2.2-6GS-SB175 |
| TRI2-6-G7 | AHB2-7GS-1na0_int2-R3 |
| TRI2-6-G9 | AHB2-9GS-1na0_int2-R3 |
| TRI2-7-G5 | AHB2-5GS-1na0_int2 |
| TRI2-7-G7 | AHB2-7GS-1na0_int2 |
| TRI2-7-G9 | AHB2-9GS-1na0_int2 |
| TRI2-8-G5 | AHB2-5GS-6msr |
| TRI2-8-G7 | AHB2-7GS-6msr |

| Abbreviation | Protein (N to C) |
|---|---|
| TRI2-3-G4 | AHB2-4GS-SB175.1 |
| TRI2-3-G6 | AHB2-6GS-SB175.1 |
| TRI2-4-G2 | AHB2-2GS-SB175.2 |
| TRI2-4-G4 | AHB2-4GS-SB175.2 |
| TRI2-4-G6 | AHB2-6GS-SB175.2 |
| TRI1-5-G2 | 36729-2GS-LCB1v2.2 |
| TRI1-5-G4 | 36729-4GS-LCB1v2.2 |
| TRI1-5-G6 | 36729-6GS-LCB1v2.2 |
| TRI1-2-G10 | SB175-10GS-LCB1v2.2 |
| TRI1-3-G6 | SB175.1-6GS-LCB1v2.2 |
| TRI1-3-G10 | SB175.1-10GS-LCB1v2.2 |
| TRI3-2-G8 | LCB3v2.2-8GS-SB175 |
| TRI3-2-G4 | LCB3v2.2-4GS-SB175 |
| FUS21-P24 | AHB2-PAS24-LCB1v2.1 |
| FUS21-P16 | AHB2-PAS16-LCB1v2.1 |
| FUS21-P12 | AHB2-PAS12-LCB1v2.2 |
| FUS21-P11 | AHB2-PAS11-LCB1v2.1 |
| FUS31-P24 | LCB3v2.2-PAS24-LCB1v2.1 |
| FUS31-P16 | LCB3v2.2-PAS16-LCB1v2.1 |
| FUS31-P11 | LCB3v2.2-PAS11-LCB1v2.1 |
| FUS23-P24 | AHB2-PAS24-LCB3v2.3 |

| | |
|---|---|
| TRI2-8-G9 | AHB2-9GS-6msr |
| TRI2-9-G5 | AHB2-5GS-1gcm |
| TRI2-9-G7 | AHB2-7GS-1gcm |
| TRI2-9-G9 | AHB2-9GS-1gcm |
| TRI2-10-G5 | AHB2-5GS-pRO-2-noHis |
| TRI2-10-G7 | AHB2-7GS-pRO-2-noHis |
| TRI2-10-G9 | AHB2-9GS-pRO-2-noHis |
| TRI2-11-G5 | AHB2-5GS-1na0_3 |
| TRI2-11-G7 | AHB2-7GS-1na0_3 |
| TRI2-11-G9 | AHB2-9GS-1na0_3 |
| TRI2-12-G5 | AHB2-5GS-4png |
| TRI2-12-G7 | AHB2-7GS-4png |
| TRI2-12-G9 | AHB2-9GS-4png |
| TRI2-2-G4 | AHB2-4GS-SB175 |
| TRI2-2-G6 | AHB2-6GS-SB175 |
| TRI2-3-G2 | AHB2-2GS-SB175.1 |

| | |
|---|---|
| FUS23-P16 | AHB2-PAS16-LCB3v2.3 |
| FUS23-P12 | AHB2-PAS12-LCB3v2.2 |
| FUS23-P11 | AHB2-PAS11-LCB3v2.3 |
| FUS32-P12 | LCB3v2.2-PAS12-AHB2v2 |
| FUS231-P24-P16 | AHB2v2-PAS24-LCB3v2.2-PAS16-LCB1v2.2 |
| FUS231-P16-P24 | AHB2v2-PAS16-LCB3v2.2-PAS24-LCB1v2.2 |
| FUS231-P16 | AHB2v2-PAS16-LCB3v2.2-PAS16-LCB1v2.2 |
| FUS231-P11-P16 | AHB2v2-PAS11-LCB3v2.2-PAS16-LCB1v2.2 |
| FUS231-P24-P11 | AHB2v2-PAS24-LCB3v2.2-PAS11-LCB1v2.2 |
| FUS231-P16-P11 | AHB2v2-PAS16-LCB3v2.2-PAS11-LCB1v2.2 |
| FUS231-P11 | AHB2v2-PAS11-LCB3v2.2-PAS11-LCB1v2.2 |
| FUS231-P7 | AHB2v2-PAS7-LCB3v2.2-PAS7-LCB1v2.2 |
| FUS321-P12 | LCB3v2.2-PAS12-AHB2v2-PAS12-LCB1v2.2 |
| FUS321-P24 | LCB3v2.2-PAS24-AHB2v2-PAS24-LCB1v2.2 |
| FUS31-G8 | LCB3v2.2-GS8-LCB1v2.2 |

**Table S2.**
Homotrimerization domains tested in this paper.

| Homotrimer # | Homotrimer Name | Reference | Notes | Protein Sequence |
|---|---|---|---|---|
| 1 | 1rfo | (*27*) | | GYIPEAPRDGQAYVRKDGEWVLLSTFL |
| 2 | SB175 | This work | Modified from SB13 (2L6HC3_13) (*84*) | SEALEELEKALRELKKSTDELERSTEELEKNPSEDALVENNRLIVENNKIIVEVLRIIAKVLK |
| 3 | SB175.1 | This work | Modified from SB175 | SPELEKALRELKKSTDELERSTEELEKNGSPEALVENNRLIVENNKIIVEVLRIIAK |
| 4 | SB175.2 | This work | Modified from SB175 | SEKALRELKKSTDELERSTEELEKNGSPEALVENNRLIVENNKIIVEVLR |
| 5 | 36729.2 | This work | Modified from 1na0_int2 | EEAELAYLLGELAYKLGEYRIAIRAYRIALKRDPNNAEAWYNLGNAYYKQGDYDEAIEYYQKALELDPNNAEAWYNLGNAYYKQGDYDEAIEYYQKALELDPNNAEAWYNLGNAYYKQGDYDEAIEYYQKALEL |
| 6 | 1na0_int2-R3 | This work | Modified from 1na0_int2 | EEAELAYLLGELAYKLGEYRIAIRAYRIALKRDPNNAEAWYNLGNAYYKQGDYDEAIEYYQKALELDPNNAEAKQNLGNAKQKQG |
| 7 | 1na0_int2 | This work | Modified from (*85*) | EEAELAYLLGELAYKLGEYRIAIRAYRIALKRDPNNAEAWYNLGNAYYKQGDYDEAIEYYQKALELDPNNAEAWYNLGNAYYKQGDYDEAIEYYQKALELDPNNAEAKQNLGNAKQKQG |
| 8 | 6msr | (*86*) | | GSEYEIRKALEELKASTAELKRATASLRASTEELKKNPSEDALVENNRLIVEHNAIIVENNRIIAAVLELIVRAIK |
| 9 | 1gcm | (*87*) | | RMKQIEDKIEEILSKIYHIENEIARIKKLIGER |
| 10 | pRO-2-noHis | (*86*) | | GSEYEIRKALEELKASTAELKRSTASLRASTEELKKNPSEDALVENNRLIVENNAIIVENNRIIAAVLELIVRAIK |
| 11 | 1na0_3 | This work | | NLAEKMYKAGNAMYRKGQYTIAIIAYTLALLKDPNNAEAWYNLGNAAYKKGEYDEAIEAYQKALELDPNNAEAWYNLGNAYYKQGDYDEAIEYYQKALELDPNNAEAKQNLGNAKQKQG |
| 12 | 4png | (*88*) | | GEIAKSLKEIAKSLKEIAWSLKEIAKSLKG |

**Table S3.**
CryoEM data collection and refinement statistics.

| | SARS-CoV-2 S/TRI2-2 EMD | SARS-CoV-2 S/TRI2-2 (local refinement) PDB EMD | SARS-CoV-2 S/FUS31-G10 (2RBD-open) EMD | SARS-CoV-2 S/FUS31-G10 (3RBD-open) EMD | SARS-CoV-2 S/FUS231-P24 (2RBD-open) EMD | SARS-CoV-2 S/FUS231-P24 (3RBD-open) EMD |
|---|---|---|---|---|---|---|
| **Data collection and processing** | | | | | | |
| Magnification | 105,000 | 105,000 | 130,000 | 130,000 | 36,000 | 36,000 |
| Voltage (kV) | 300 | 300 | 300 | 300 | 200 | 200 |
| Electron exposure ($e^{-}$/Å$^2$) | 60 | 60 | 70 | 70 | 60 | 60 |
| Defocus range (μm) | -0.5 – -2.5 | -0.5 – -2.5 | -0.5 – -2.5 | -0.5 – -2.5 | -0.5 – -2.5 | -0.5 – -2.5 |
| Pixel size (Å) | 0.4215 | 0.4215 | 0.525 | 0.525 | 1.16 | 1.16 |
| Symmetry imposed | C1 | C1 | C1 | C1 | C1 | C1 |
| Final particle images (no.) | 75,519 | 206,541 | 9,733 | 10,649 | 112,075 | 18,084 |
| Map resolution (Å) | 2.8 | 2.9 | 4.57 | 4.65 | 3.9 | 5.2 |
| FSC threshold | 0.143 | 0.143 | 0.143 | 0.143 | 0.143 | 0.143 |
| Map sharpening *B*factor (Å$^2$) | -63 | -31 | -63 | -71 | -99 | -167 |
| **Validation** | | | | | | |
| MolProbity score | | 1.03 | | | | |
| Clashscore | | 1.62 | | | | |
| Poor rotamers (%) | | 0 | | | | |
| Ramachandran plot | | | | | | |
| Favored (%) | | 97.37 | | | | |
| Allowed (%) | | 2.25 | | | | |
| Disallowed (%) | | 0.38 | | | | |

**Table S4.**
Table of DNA and protein sequences for multivalent minibinders used in this manuscript. DNA sequences are the open reading frame coding for the expressed protein. Protein sequences are annotated as follows. Minibinder and homotrimer sequences are denoted by square brackets []. Secondary sequences (e.g., expression tag, purification tag, etc.) are annotated by parenthesis (). Non-minibinder or non-homotrimer sequences are annotated by curly brackets {} (captures linkers and other secondary sequences). (Separate csv file)
